# HLA-DRB1*15:01 drives sex- and age-dependent microglial activation and neuroimmune signaling

**DOI:** 10.64898/2026.03.11.711132

**Authors:** EM Reyes-Reyes, D Chinnasamy, F Fernandez, MD Trial, VD Nguyen, Q He, C Figueroa, AC Leslie, D Bradford, JP Wiegand, KE Rodgers

## Abstract

**Introduction:** The major histocompatibility complex class II (MHC-II) pathway is central to adaptive immunity and immune tolerance, and age-related erosion of these mechanisms is increasingly recognized as a driver of chronic neuroinflammation. The HLA-DRB1*15:01 allele—the strongest genetic risk factor for multiple sclerosis in Caucasians—has been implicated in shaping pathogenic CD4⁺ T-cell responses and broader neuroimmune vulnerability, yet how this allele modulates age- and sex-dependent neuroimmune processes within the central nervous system (CNS) remains poorly defined.

**Methods:** We investigated the impact of HLA-DRB1*15:01 expression using a humanized mouse model (HLA mice) and wild-type (WT) controls. Male and female mice were analyzed at 6, 9, and 15 months of age, with endocrine stratification in females. Behavioral testing, flow cytometry, immunofluorescence, and multiplex cytokine analyses were used to assess cognitive performance, glial activation and oxidative stress, astrocyte–microglia IL-3/IL-3R signaling, endothelial activation, selective immune cell accumulation at CNS borders, tissue organization, and hippocampal cytokine profiles.

**Results:** HLA mice developed age- and sex-dependent cognitive impairment, most pronounced in aged females. HLA-DRB1*15:01 expression promoted progressive microglial activation, characterized by increased CD14 and CD68 expression, elevated mitochondrial oxidative stress, altered astrocyte phenotypes, and enhanced IL-3/IL-3R signaling. Hippocampal axonal and myelin organization was disrupted in aged HLA mice, and this disruption was spatially associated with increased microglial presence. At CNS interfaces, HLA mice exhibited selective immune remodeling, including increased accumulation of CD4⁺ T cells and NK1.1⁺CD3⁺ natural killer T (NKT) cells, particularly in females, accompanied by endothelial activation, as evidenced by elevated ICAM-1 and E-selectin expression. Hippocampal cytokine profiling revealed selective, sex-biased alterations, including increased IL-12p70 and reduced IL-10 and IL-2, without broad induction of classical inflammatory cytokines.

**Conclusion:** Together, these findings demonstrate that HLA-DRB1*15:01 drives a coordinated, age- and sex-dependent neuroinflammatory program linking behavioral dysfunction, glial activation and oxidative stress, selective immune cell recruitment, endothelial activation, tissue remodeling, and targeted cytokine imbalance. This integrated phenotype provides mechanistic insight into how this major MS risk allele confers vulnerability to chronic neuroinflammation during aging, with heightened impact in females.

## 1 Introduction

The major histocompatibility complex class II (MHC-II) pathway plays a central role in adaptive immunity by enabling antigen-presenting cells (APCs) to present extracellular peptides to CD4⁺ T lymphocytes. During early life, presentation of self-antigens plays a critical role in establishing central and peripheral immune tolerance by eliminating or functionally inactivating autoreactive T cells through thymic deletion, induction of anergy, and suppression by regulatory T cells. With aging, however, tolerance mechanisms progressively decline, driven by reduced regulatory T-cell function, altered sensitivity of effector T cells to suppressive signals, and cumulative inflammatory exposure. This age-associated erosion of immune tolerance may permit the activation of previously restrained autoreactive T cells, thereby promoting chronic, low-grade neuroinflammation (1, 2).

Human leukocyte antigen (HLA) class II molecules—including HLA-DR, HLA-DQ, and HLA-DP—are highly polymorphic, and variation within these loci represents a major genetic determinant of autoimmune disease susceptibility. Among these, polymorphisms in HLA-DRB1 have been most consistently linked to immune-mediated disorders, underscoring the importance of antigen presentation in shaping pathogenic CD4⁺ T-cell responses (3–5). The HLA-DRB1*15:01 allele is the strongest and most reproducible genetic risk factor for multiple sclerosis (MS), conferring approximately a 3-fold increase in disease susceptibility (6). Beyond MS, HLA-DRB1*15:01 has been associated with several other autoimmune conditions, including juvenile idiopathic arthritis, Sjögren’s syndrome, and systemic lupus erythematosus. Many of these disorders exhibit cognitive dysfunction or central nervous system (CNS) involvement, suggesting that HLA-DRB1*15:01–associated immune dysregulation is not restricted to peripheral autoimmunity (7–13). Moreover, emerging genetic and epidemiological studies implicate variation within the HLA-DRB1*15-01 haplotype in susceptibility to late-onset neurodegenerative diseases, including Alzheimer’s disease (14–16) and Parkinson’s disease (17, 18), pointing to a broader role for this allele in age-related neuroimmune vulnerability. Mechanistically, HLA-DRB1*15:01 shapes the repertoire, activation state, and cytokine polarization of CD4⁺ T cells through selective presentation of self and foreign peptides(17, 19, 20). In MS lesions, HLA-DRB1*15:01 molecules have been shown to present myelin basic protein (MBP)–derived epitopes in situ, and structural studies demonstrate stable binding of the immunodominant MBP_85-99_ peptide within the HLA-DRB1*15:01 peptide-binding groove (21, 22). HLA-DRB1*15:01 can also bind peptides derived from proteins associated with neurodegeneration, including amyloid-β (15) and α-synuclein (17), suggesting that this allele contributes to immune recognition of neuronal and glial self-antigens under permissive inflammatory conditions. These findings provide direct molecular evidence linking this allele to autoreactive CD4⁺ T-cell responses within the CNS. Despite the well-established role of HLA-DRB1*15:01 in shaping CD4⁺ T-cell responses (17, 19, 20), how this allele influences age- and sex-dependent neuroimmune interactions within the CNS remains poorly defined. In particular, it is unclear how HLA-DRB1*15:01 expression modulates resident glial states, neuroimmune signaling pathways, immune–CNS interfaces, and tissue-level organization during aging in the absence of overt autoimmune challenge. To isolate HLA-DRB1*15:01–specific mechanisms in vivo, we used a previously established transgenic mouse model generated on an MHC-II–null background that expresses the human HLA-DRB1*15:01 peptide-binding domains under the control of the murine I-E promoter (22). This design ensures physiological, antigen-presenting cell–restricted expression of HLA-DRB1*15:01 and supports normal CD4⁺ T-cell development and function. These mice develop chronic experimental autoimmune encephalomyelitis (EAE) following immunization with myelin oligodendrocyte glycoprotein, establishing that HLA-DRB1*15:01–restricted CD4⁺ T cells are sufficient to drive CNS autoimmunity in vivo (22). However, the impact of HLA-DRB115:01 expression on spontaneous, age-related neuroinflammatory processes, independent of induced autoimmunity, has not been systematically examined. In the present study, we investigated how HLA-DRB1*15:01 expression shapes neuroimmune aging by integrating behavioral, cellular, molecular, and tissue-level analyses in male and female mice across multiple ages. We examined microglial oxidative stress and activation, astrocyte–microglia cytokine signaling, endothelial activation, immune cell accumulation at CNS interfaces, hippocampal cytokine profiles, and associated behavioral outcomes. Together, these studies define how a major human autoimmune risk allele intersects with aging and sex to remodel the neuroimmune environment, providing mechanistic insight into how genetic susceptibility may promote chronic neuroinflammation and CNS vulnerability over the lifespan.

## 2 Material and methods

### 2.1 Animals and brain cells isolation

All animal studies and procedures were conducted in accordance with the National Institutes of Health guidelines for the care and use of laboratory animals and were approved by the University of Arizona Institutional Animal Care and Use Committee (IACUC). Female and male HLA-DRB1*15:01 transgenic mice were obtained from Dr. Vandenbark; the generation and characterization of this mouse strain have been described previously. Age-matched B6D2F1/J wild-type mice were obtained from The Jackson Laboratory (JAX:100006) at 12 weeks of age. All mice were bred and/or aged in a health-designated (HD) animal facility under specific-pathogen-free conditions (Helicobacter- and murine norovirus–negative), with non-sterile food and housing. Mice were anesthetized with isoflurane at 6, 9, or 15 months of age, and blood was collected by cardiac puncture. Animals were then perfused transcardially with cold phosphate-buffered saline (PBS), and brains were harvested. Brain tissue was dissociated into single-cell suspensions using the Miltenyi Biotec Adult Brain Dissociation Kit (Cat. no. 130-107-677) according to the manufacturer’s instructions.

### 2.2 Nesting test

Nestlet shredding and nest construction were assessed as measures of spontaneous, goal-directed home-cage behavior. Mice were individually housed for 24 hours in clean cages containing ∼1 cm of Alpha-Dri bedding and provided with a single 5 × 5 cm compressed cotton Nestlet (Ancare, USA), with ad libitum access to food and water. Cages were labeled to blind the experimenter to genotype and experimental group. After 18 hours, cages were inspected and overhead photographs were obtained. Nesting behavior was scored using a standardized 1–5 scale adapted from established protocols, in which scores reflected the extent of Nestlet shredding and nest structure: 1 = Nestlet largely intact (>90% intact); 2–3 = partial to extensive shredding without a consolidated nest site; and 4–5 = formation of an identifiable to near-perfect crater-shaped nest with shredded material gathered into a defined structure. Scoring was performed by an experimenter blinded to experimental conditions.

### 2.3 Novel object recognition (NOR) test

Intermediate memory was assessed using the novel object recognition (NOR) test. Mice were first given a 24-hour habituation and familiarization period to acclimate to the testing environment and a set of identical familiar objects. NOR testing was conducted in arenas measuring 32 × 26 × 30 cm, with up to eight arenas prepared simultaneously. Two identical familiar objects were placed diagonally within each arena, equidistant from the arena walls. During the familiarization phase, each mouse was allowed to freely explore the arena and the two familiar objects for 15 minutes. Twenty-four hours later, the test phase was conducted by replacing one of the familiar objects with a novel object positioned in the same location and orientation as the original familiar object. Mice were again allowed to explore the arena for 15 minutes. Behavior during the test phase was recorded using an overhead camera and analyzed with an automated tracking system (AnyMaze software, Stoelting Co.). The software tracked each mouse throughout the test session and quantified the time spent exploring the novel and familiar objects, defined as directed head orientation toward the object. Exploration times were used to calculate the following indices: Recognition Index (%) = 100 × [novel object exploration time / (novel + familiar object exploration time)] and Discrimination Index (%) = 100 × [(novel object exploration time – familiar object exploration time) / (novel + familiar object exploration time)]

### 2.4 Immunohistochemistry

Immunohistochemistry was performed as previously described (4). Briefly, brain hemispheres were fixed overnight in 4% paraformaldehyde at 4 °C and subsequently transferred to 30% sucrose for cryoprotection. Tissue was embedded in optimal cutting temperature (OCT) compound and sectioned using a cryostat. Sections were blocked for 60 minutes at room temperature in 0.5% Triton X-100 in PBS containing 5% normal goat serum (NGS). Primary antibodies (Supplemental Table 1) were diluted in 0.5% Triton X-100/PBS with 5% NGS and applied overnight at 4 °C. Sections were then thoroughly washed in 0.5% Triton X-100/PBS and incubated for 1 hour at room temperature with the appropriate secondary antibodies (Table 1), diluted in the same blocking solution. Following washing, sections were mounted onto glass slides using a DAPI-containing mounting medium and coverslipped. Immunofluorescent images were acquired using a Zeiss LSM 880 NLO confocal microscope at 20× magnification.

### 2.5 Flow Cytometry Staining

Single-cell suspensions prepared from mouse brain tissue were used for multiparameter flow cytometry analyses. All staining procedures were performed using ice-cold buffers unless otherwise indicated, and samples were protected from light throughout. Prior to staining with cell-surface marker antibodies, cells were incubated with an Fc receptor–blocking reagent for 15 minutes at 4 °C to prevent nonspecific antibody binding. For assessment of oxidative stress, cells were incubated with 5 µM MitoSOX™ Red (Thermo Fisher Scientific, #M36008) in DPBS supplemented with glucose at 37 °C for 20 minutes in the absence of CO₂. Cells were then stained with cell-surface marker antibodies for 25 minutes at 4 °C, washed with PBS, and centrifuged at 400 × g for 5 minutes. For intracellular staining, cells were fixed using eBioscience™ IC Fixation Buffer (Thermo Fisher Scientific, #00-8222-49) for 15 minutes, followed by permeabilization with eBioscience™ Permeabilization Buffer (Thermo Fisher Scientific, #00-8333-56) for 20 minutes. After washing, cells were incubated with antibodies recognizing intracellular proteins for 25 minutes at room temperature. Cell pellets were resuspended in PBS for acquisition. For unfixed samples, DAPI was added immediately prior to data acquisition to exclude dead cells. Samples were analyzed by flow cytometry using a MACSQuant 10 (Miltenyi Biotec), and data were analyzed with FlowLogic software.

### 2.6 Flow cytometry cell population definitions

Antibody combinations were selected to enable accurate identification of specific immune and glial cell populations and their functional states based on established marker expression. Antibody details are provided in Supplemental Table 1. Microglia were defined as CD45 ^low/int^ CD11b⁺ Ly-6C⁻ cells, where CD45 was used to identify leukocytes, CD11b identified the myeloid lineage, and Ly-6C was used to exclude infiltrating monocytes. Within the microglial population, activation- and stress-associated markers were quantified, including CD14 (innate immune activation–associated receptor), CD68 (lysosomal/phagolysosomal marker indicative of microglial activation), CD123 (IL-3Rα, IL-3 receptor expression), and MitoSOX™ Red (mitochondrial superoxide production). Astrocytes were identified by surface expression of ACSA-2. Intracellular staining was used to assess astrocytic functional markers, including IL-3, an astrocyte-derived cytokine, and GFAP, an intermediate filament protein associated with astrocytic structure and reactivity. T-cell populations were identified using CD45 to define leukocytes and CD3ε to identify T-cell lineage. T-cell subsets were further distinguished by expression of CD4 (helper T cells) and CD8β (cytotoxic T cells). NK1.1 expression in combination with CD3ε was used to identify NK1.1⁺CD3ε⁺ natural killer T (NKT) cells. Endothelial cells were identified by surface expression of CD31 (PECAM-1), a pan-endothelial marker. Endothelial activation status was assessed by measuring the expression of adhesion molecules involved in leukocyte tethering, adhesion, and transmigration, including CD54 (ICAM-1), CD106 (VCAM-1), CD62E (E-selectin), and CD62P (P-selectin). ICAM-1 and VCAM-1 were evaluated as indicators of firm leukocyte adhesion, whereas E-selectin and P-selectin were assessed as markers of endothelial activation associated with leukocyte rolling and recruitment at the neurovascular interface.

### 2.7 Flow cytometry gating strategy

The sequential gating strategies used to identify these populations using Flow Logic software are illustrated in Supplementary Figures 1–5. For all panels, cells were first gated on forward scatter (FSC-A) versus side scatter (SSC-A) to exclude debris. Dead cells were excluded using DAPI as indicated. Doublets were removed using FSC-A versus FSC-H gating.

#### Panel 1: Microglia identification and mitochondrial superoxide

Microglia were identified as CD11b⁺ CD45^low/int^ cells, with peripheral myeloid cells excluded based on Ly6C expression. Mitochondrial superoxide production was assessed using MitoSOX-PE, with fluorescence quantified within the gated microglial population (Supplementary Figure 1).

#### Panel 2: Microglial activation

Using the same microglial gating strategy for microglia identification, activation status was evaluated by analysis of CD14 and CD68 expression within CD11b⁺ CD45^low/int^ Ly6C⁻ microglia (Supplementary Figure 2).

#### Panel 3: IL-3 and IL-3R in astrocytes and microglia

Following exclusion of debris and doublets, CD11b and CD45 expression were used to separate myeloid (CD11b⁺ CD45^low/int^) from non-myeloid (CD11b⁻ CD45⁻) populations. Astrocytes were identified within the CD11b⁻ CD45⁻ fraction based on ACSA-2 expression, and IL-3 and GFAP expression were assessed within the ACSA-2⁺ astrocyte population. In parallel, microglia were defined as CD11b⁺ CD45^low/int^ Ly6C⁻ ACSA-2⁻ cells, and IL-3R expression was quantified within this gated microglial population (Supplementary Figure 3).

#### Panel 4: T cell and NKT cells

Following exclusion of debris, dead cells, and doublets, T cells were defined as CD3⁺ CD45⁺ cells. Within this population, NKT cells were identified as NK1.1⁺ CD3⁺ CD45⁺ cells. T cells were further subdivided into CD4⁺ CD3⁺ CD45⁺ NK1.1⁻ and CD8⁺ CD3⁺ CD45⁺ NK1.1⁻ subsets (Supplementary Figure 4).

#### Panel 5: Endothelial cell activation

Endothelial cells were identified from brain single-cell suspensions following exclusion of debris, dead cells, and doublets. Endothelial cells were defined as CD31⁺ cells, and activation status was assessed by expression of ICAM-1, VCAM-1, E-selectin (CD62E), and P-selectin (CD62P) within the CD31⁺ population (Supplementary Figure 5).

### 2.8 Hippocampus homogenization

Frozen tissue samples were homogenized to extract total protein. Approximately 15-25mg of tissue was placed into a 1.5mL microcentrifuge tube (VWR North American Cat. No. 20170-038) containing 350μL of lysis buffer (50mM Tris HCl pH 7.5, 150mM NaCl, 2mM Na2EDTA, 1% NP-40, 0.25% deoxycholate) supplemented with protease (Millipore Sigma Cat. No. 539134) and phosphatase (Millipore Sigma Cat. No. 524629) inhibitors. Four to five 0.5mm silica glass beads were then added to the samples before being mechanically homogenized using the Next Advance Bullet Blender® GOLD for 4 minutes at a speed setting of 4. The silica glass beads were subsequently removed from the homogenized solution before being centrifuged at 4°C 22,000 xg for 30 minutes. After centrifugation, sample supernatant was carefully collected and aliquoted before being frozen at -80°C.

### 2.9 Assessment of proinflammatory cytokines

Proinflammatory cytokines in hippocampal homogenates were quantified using the V-Plex Proinflammatory Panel 1 Mouse Kit (Meso Scale Discovery Cat. No. K15048D-1). Total protein concentrations of all individual samples were measured through the BCA Protein Assay (Thermo Scientific Cat. No. 23228) and normalized to a concentration of 2mg/mL prior to sample loading. 25μL of normalized sample was loaded in duplicate into each well and all subsequent steps were performed according to the manufacturer’s instructions.

### 2.10 Statistical analysis

Data were analyzed using GraphPad Prism (GraphPad Software, San Diego, CA, USA). Datasets were evaluated using three-way analysis of variance (ANOVA) to assess the effects of genotype, age, and sex, followed by Bonferroni post hoc correction for multiple comparisons where applicable. Statistical significance was set at p < 0.05. Data are presented as mean ± standard deviation (SD).

## 3 RESULTS

### 3.1 HLA-DRB1*15:01 expression leads to age- and sex-dependent behavioral impairments

To determine whether expression of the human MS risk allele HLA-DRB1*15:01 is associated with age-dependent neurobiological dysfunction, wild-type (WT) and humanized HLA mice were assessed at 6, 9, and 15 months of age using behavioral assays that serve as functional readouts of brain integrity. These included measures of daily living, goal-directed behavior (nest building, which is known to be impaired in several neurological disorders and diseases), and cognitive function, specifically recognition memory (novel object recognition) and spatial memory (object-location discrimination) (23–26).

Behavioral performance was preserved in WT mice of both sexes at all ages and in HLA male mice across all ages examined. In contrast, behavioral impairments were observed selectively in 15-month-old HLA female mice, while 6- and 9-month-old HLA females showed no deficits. Nest-building behavior (**Figure 1A**) was significantly impaired in 15-month-old HLA female mice, which exhibited a 49% reduction in nest construction compared with age-matched WT female mice (p < 0.0001). Nesting scores in 15-month-old HLA female mice were also significantly lower than those of 6-month-old HLA female mice (p = 0.006) and 9-month-old HLA female mice (p < 0.0001). Deficits in daily living behavior were accompanied by impairments in recognition memory, as assessed using the novel object recognition (NOR) task (**Figure 1B**). 15-month-old HLA female mice displayed a 40% reduction in recognition index compared with age-matched WT female mice (p = 0.0043). Recognition index values in 15-month-old HLA female mice were also significantly reduced compared with 6-month-old HLA female mice (p < 0.0001) and 9-month-old HLA female mice (p = 0.019). Analysis of object-location discrimination (**Figure 1C**) revealed a similar pattern. 15-month-old HLA female mice exhibited negative discrimination indices, indicating impaired spatial recognition. These values were significantly reduced compared with age-matched WT female mice (p= 0.0039), 6-month-old HLA female mice (p < 0.0001), and 9-month-old HLA female mice (p= 0.029).

**Figure 1.**
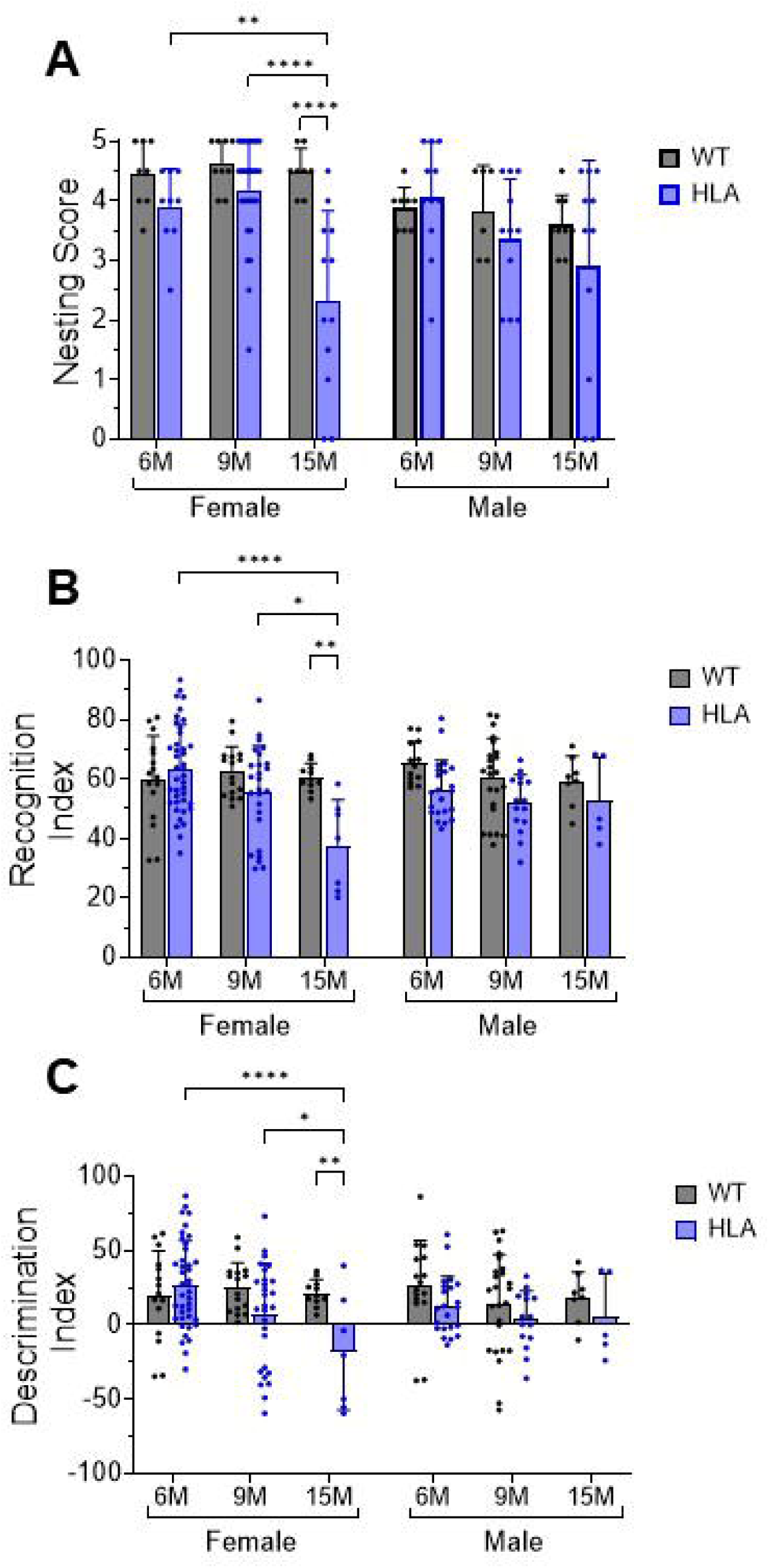
Behavioral assessment of wild-type and HLA-DRB1*15:01 mice across age and sex. Wild-type (WT, gray) and HLA-DRB1*15:01 (HLA, blue) mice were tested at 6, 9, and 15 months of age. (A) Nest-building score, (B) novel object recognition (NOR) index, and (C) object-location discrimination. Bars represent mean ± SD with individual data points shown; n = 10–24 mice per group. Statistical analyses were performed using two-way or three-way ANOVA as appropriate, followed by Bonferroni’s multiple-comparisons test. *p < 0.05, **p < 0.01, ****p < 0.0001.

Together, these findings identify a late-onset behavioral phenotype restricted to female HLA mice, providing functional evidence that HLA-DRB1*15:01 expression is associated with increased vulnerability of the aging female brain. This behavioral profile motivated subsequent analyses of the cellular and molecular neuroimmune mechanisms underlying this phenotype.

### 3.2 HLA-DRB1*15:01 expression is associated with age- and sex-dependent increases in microglial oxidative stress

Mitochondrial oxidative stress in microglia is an indicator of neuroimmune dysfunction and has been implicated in age-related neuroinflammatory and neurodegenerative processes (27, 28). We therefore examined whether the behavioral vulnerability observed in HLA mice was accompanied by age- and sex-dependent alterations in microglial oxidative state.

Mitochondrial oxidative stress in brain microglia was evaluated using MitoSOX fluorescence as a readout of mitochondrial oxidant burden (29). Representative flow cytometry dot plots are shown in Figure 2a, with quantitative analysis presented in Figure 2b. In HLA female mice, the proportion of MitoSOX-positive microglia increased from approximately 1.5% at 6 months to ∼11.5% at 9 months, reaching ∼21% at 15 months. In HLA male mice, MitoSOX-positive microglia increased from approximately 1.0% at 6 months to ∼11.5% at 9 months, reaching ∼13.5% at 15 months. In contrast, WT mice of both sexes displayed consistently low frequencies of MitoSOX-positive microglia across all ages examined, remaining below 5% (**Figure 2A, B**).

**Figure 2.**
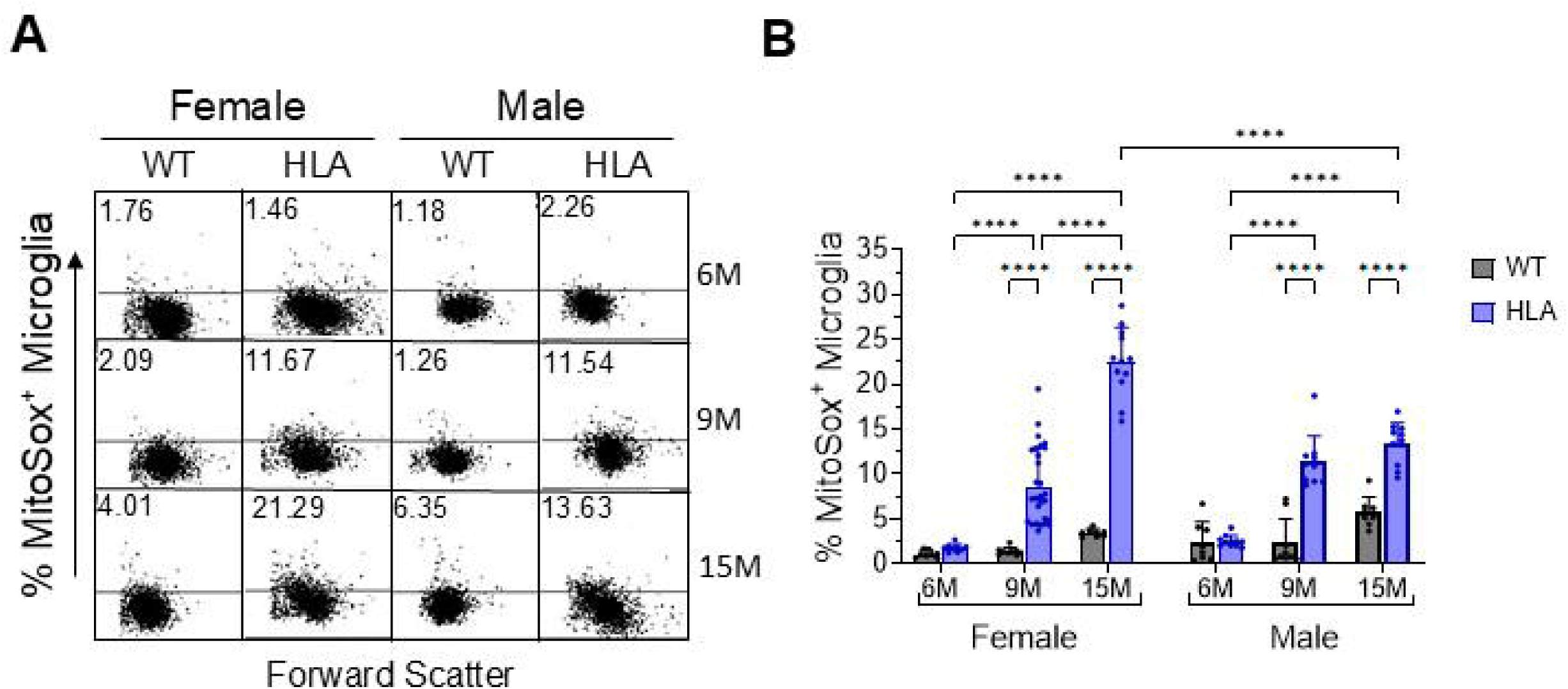
Microglial mitochondrial oxidative stress across age and sex in wild-type and HLA-DRB1*15:01 mice. Brain cells from female and male wild-type (WT, gray) and HLA-DRB1*15:01 (HLA, blue) mice at 6, 9, and 15 months of age (6M, 9M, 15M) were dissociated, stained with cell type–specific markers, and analyzed by flow cytometry. Microglia were identified as Ly6C⁻ CD11b^high CD45^int cells. Mitochondrial oxidative stress was evaluated using MitoSOX fluorescence as a readout of mitochondrial oxidant burden, and data are expressed as the percentage of MitoSOX⁺ microglia. (A) Representative flow cytometry dot plots. (B) Quantitative analysis. Bars represent mean ± SD, with individual mice shown (n = 8–18 per group). Data were analyzed by three-way ANOVA (Age × Sex × Genotype) followed by Bonferroni’s multiple-comparisons test.. *p < 0.05, **p < 0.01, ****p < 0.0001. Significant main effects of age (p < 0.0001) and genotype (p < 0.0001) were detected. Significant Age × Sex, Age × Genotype, and Age × Sex × Genotype interactions (all p < 0.0001), as well as a significant Sex × Genotype interaction (p = 0.0011).

Post hoc analyses demonstrated that HLA mice of both sexes exhibited significantly higher frequencies of MitoSOX-positive microglia than age-matched WT mice at 9 and 15 months (e.g., 15-month WT male vs 15-month HLA male, p < 0.0001), whereas no significant genotype differences were detected at 6 months. Notably, at 15 months of age, the frequency of MitoSOX-positive microglia was higher in HLA females than in HLA males, indicating a sex-dependent divergence in microglial oxidative stress at advanced age **(Figure 2B)**.

Together, these findings indicate that expression of HLA-DRB1*15:01 is associated with early, age- and sex-dependent alterations in microglial oxidative state, which emerge prior to overt inflammatory activation. The selective amplification of oxidative stress in aged HLA females parallels the late-onset behavioral vulnerability observed in this group, suggesting a potential cellular context in which downstream neuroimmune changes may occur. These observations prompted further examination of whether increased oxidative burden is associated with alterations in microglial activation phenotype.

### 3.3 HLA-DRB1*15:01 expression is associated with age- and sex-dependent microglial activation in the brain

Microglial activation is a central cellular feature of neuroinflammatory and neurodegenerative processes(28, 30). We examined whether the age-dependent increase in microglial oxidative stress was accompanied by changes in microglial activation state. MHC class II expression was not used as a marker of microglial activation in this study, as the HLA-DRB1*15:01 humanized mouse model lacks endogenous murine MHC class II expression. Accordingly, microglial activation was evaluated using CD14, a pattern-recognition receptor associated with innate immune and inflammatory signaling (31–33), and CD68, a lysosomal/phagolysosomal marker associated with microglial activation and phagocytic activity (34). These markers provide MHC-II–independent measures suitable for comparative analyses between WT and HLA mice.

Brain cells were isolated from wild-type (WT) and HLA-DRB1*15:01 (HLA) mice at 6, 9, and 15 months of age, in both females and males, and analyzed by flow cytometry (**Figure 3**). In HLA female mice, the proportion of CD14⁺ microglia increased progressively with age, rising from <0.5% at 6 months to approximately 2% at 9 months, and exceeding 5% at 15 months. WT females also showed an age-associated increase, with significantly higher CD14⁺ microglia at 15 months compared with 6 months, but levels remained substantially lower than those observed in age-matched HLA females. Post hoc analyses confirmed that HLA females exhibited significantly higher frequencies of CD14⁺ microglia than WT females at 15 months (**Figure 3A, B**). In HLA male mice, CD14⁺ microglia remained below 0.5% at 6 and 9 months, but increased to approximately 2.0 % at 15 months, whereas WT males showed no significant age-dependent increase. As a result, a sex difference was evident at 15 months, with higher frequencies of CD14⁺ microglia in HLA females (∼5%) than in HLA males (∼2%) (**Figure 3A, B**).

**Figure 3.**
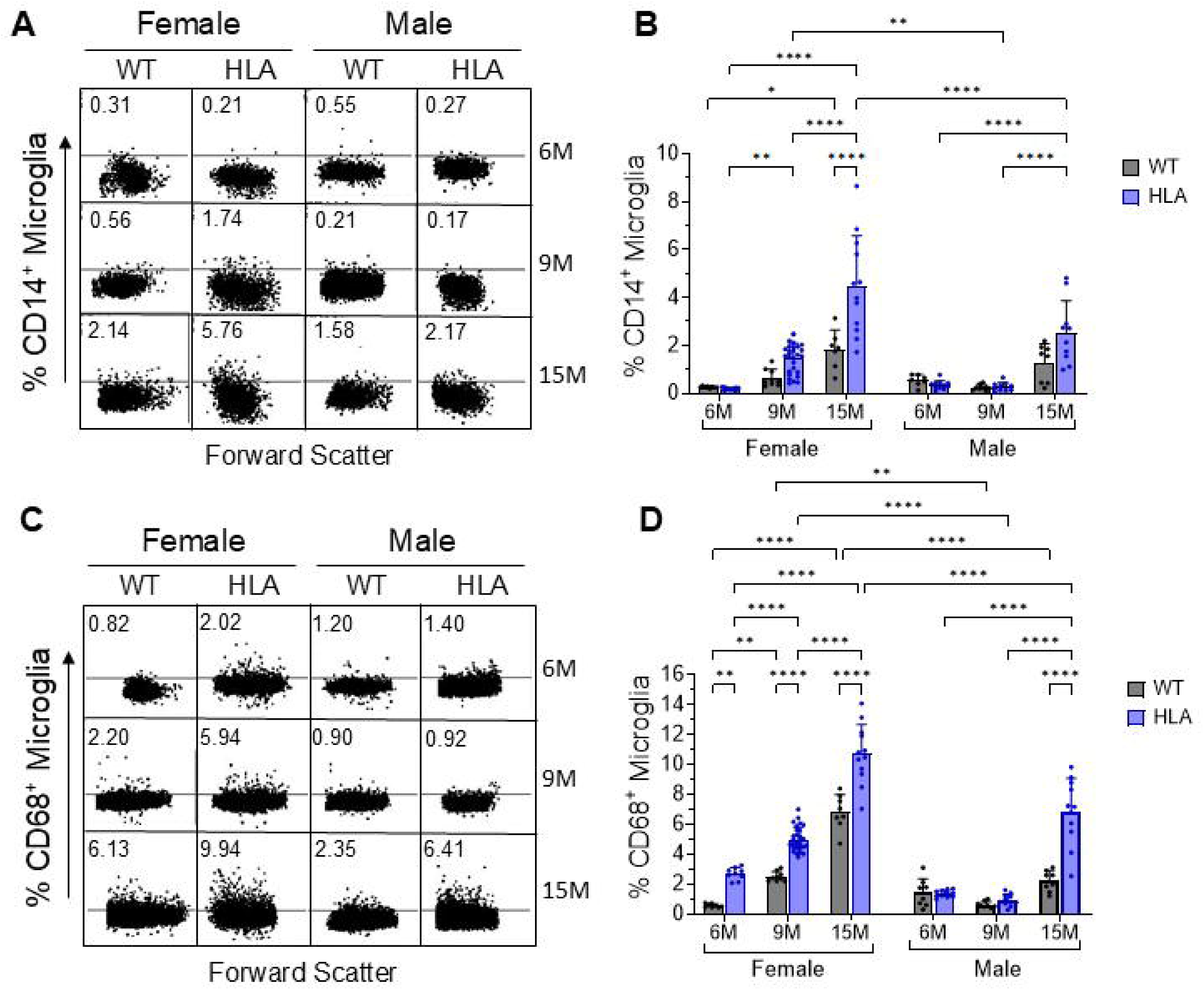
Age- and sex-dependent microglial activation in wild-type and HLA-DRB1*15:01 mice. Brain cells were dissociated from female and male wild-type (WT, gray) and HLA-DRB1*15:01 (HLA, blue) mice at 6, 9, and 15 months of age, stained with cell type–specific markers, and analyzed by flow cytometry. Cells were gated based on forward- and side-scatter characteristics, with dead cells and doublets excluded. Microglia were identified as Ly6C⁻ CD11b^^high^ CD45^^int^ cells. (A) Representative flow cytometry dot plots and (B) quantification of CD14⁺ microglia, a pattern-recognition receptor associated with innate immune and inflammatory signaling. (C) Representative flow cytometry dot plots and (D) quantification of CD68⁺ microglia, a lysosomal/phagolysosomal marker of microglial activation. Bars represent mean ± SD, with individual mice shown (n = 8–18 per group). Data were analyzed by three-way ANOVA (Age × Sex × Genotype) followed by Bonferroni’s multiple-comparisons test. *p < 0.05, **p < 0.01, ****p < 0.0001. For CD14⁺ microglia, significant main effects of age (p < 0.0001), sex (p = 0.0002), and genotype (p < 0.0001) were detected, along with significant Age × Sex (p = 0.0013), Age × Genotype (p < 0.0001), and Sex × Genotype (p = 0.0157) interactions; the Age × Sex × Genotype interaction was not significant (p = 0.2797). For CD68⁺ microglia, three-way ANOVA revealed significant main effects of age, sex, and genotype (all p < 0.0001), as well as significant Age × Sex, Age × Genotype, Sex × Genotype (p = 0.0024), and Age × Sex × Genotype (p = 0.0022) interactions.

We next assessed CD68 expression to evaluate phagolysosomal microglial activation. In female mice, the frequency of CD68⁺ microglia increased with age in both genotypes. WT females showed an increase from approximately 1% at 6 months to ∼2% at 9 months and ∼6% at 15 months, whereas HLA females exhibited a larger increase, from approximately 2 % at 6 months to ∼6% at 9 months and ∼10% at 15 months (**Figure 3C, D**). Post hoc analyses demonstrated that HLA females had significantly higher frequencies of CD68⁺ microglia than age-matched WT females at 6 months (p = 0.0012), 9 months (p < 0.0001), and 15 months (p < 0.0001) (**Figure 3D**). In male mice, WT animals showed no significant age-dependent changes in CD68⁺ microglia (< 2.5%). In contrast, HLA males showed CD68 expression levels of ∼1–1.5% at 6 and 9 months, but exhibited a marked increase at 15 months to ∼6.5%, which was significantly higher than those of age-matched WT males (p < 0.0001) (**Figure 3C, D**). Consistent with the significant Sex × Genotype and Age × Sex × Genotype interactions, sex-dependent differences in CD68⁺ microglia were evident at both 9 and 15 months, with higher frequencies in HLA females (∼6% vs ∼1% at 9 months; ∼10% vs ∼6.5% at 15 months) compared with HLA males (**Figure 3C, D**).

These data show that HLA-DRB1*15:01 expression is associated with progressive, age- and sex-dependent microglial activation, encompassing both innate immune signaling and phagolysosomal pathways. The greater magnitude of activation observed in aged HLA females suggests that microglial responses to aging are modulated by sex in the context of HLA expression. Given the central role of activated microglia in shaping the neuroinflammatory environment, these findings raised the possibility that intercellular glial communication may also be altered in HLA mice.

### 3.4 HLA-DRB1*15:01 promotes age- and sex-dependent astrocyte–microglia immune signaling

Astrocytes and microglia actively participate in neuroinflammation by regulating the innate immune system. Given the emergence of age- and sex-dependent behavioral impairments, increased microglial oxidative stress, and changes in activation-associated microglial markers in mice expressing HLA-DRB1*15:01, we next examined whether these phenotypes were accompanied by alterations in glial immune signaling pathways that regulate microglial activation state. Because astrocyte–microglia communication plays a central role in shaping neuroinflammatory responses, we focused on the IL-3/IL-3 receptor axis, which rather than serving as a direct marker of activation. We therefore quantified age-, sex-, and genotype-dependent changes in IL-3R expression on microglia, IL-3 production by astrocytes, and astrocyte reactivity to assess coordinated glial immune signaling in WT and HLA mice.

IL-3R⁺ microglia increased robustly with age in both genotypes, with consistently higher frequencies in females than in males (**Figure 4A, B**). In WT females, IL-3R⁺ microglia increased from ∼9% at 6 months to ∼19% at 9 months and ∼21% at 15 months, whereas HLA females showed a markedly greater increase, rising from ∼16% at 6 months to ∼32% at 9 months and ∼42% at 15 months. HLA females exhibited significantly higher IL-3R⁺ microglial frequencies than age-matched WT females at all ages examined (all p < 0.0001). In males, WT mice showed relatively stable IL-3R⁺ microglial frequencies (∼10%) at 6 and 9 months, followed by an increase to ∼20% at 15 months. HLA males exhibited a significant age-dependent increase, rising from ∼11% at 6 months to ∼28% at 9 months and ∼33% at 15 months, with levels at 15 months significantly higher than those of age-matched WT males (p < 0.0001) (**Figure 4B**). Across both genotypes, females exhibited higher IL-3R⁺ microglial frequencies than males, indicating a sex bias that became more pronounced with aging.

**Figure 4.**
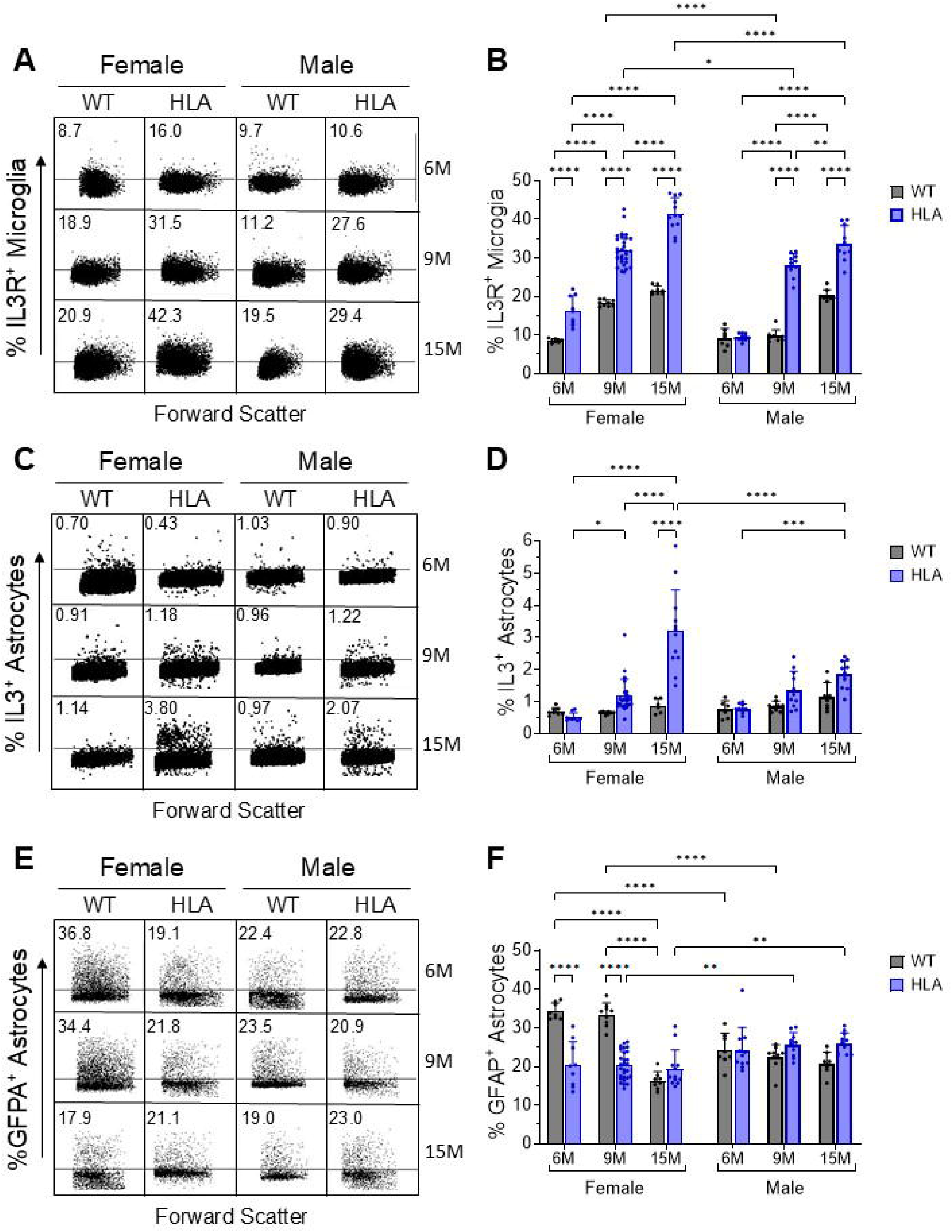
Age- and sex-dependent modulation of astrocyte–microglia IL-3/IL-3R signaling by HLA-DRB1*15:01. Brain cells were dissociated from female and male wild-type (WT, gray) and HLA-DRB1*15:01 (HLA, blue) mice at 6, 9, and 15 months of age, stained with cell type–specific markers, and analyzed by flow cytometry. Cells were gated based on forward- and side-scatter characteristics, with dead cells and doublets excluded using DAPI staining and FSC-A/FSC-H parameters. Microglia were identified as Ly6C⁻ CD11b^^high^ CD45^^int^ cells, and astrocytes were identified by ACSA-2 expression. (A) Representative flow cytometry dot plots and (B) quantification of IL-3R⁺ microglia. (C) Representative flow cytometry dot plots and (D) quantification of IL-3⁺ astrocytes. (E) Representative flow cytometry dot plots and (F) quantification of GFAP⁺ astrocytes. Bars represent mean ± SD, with individual mice shown (n = 8–18 per group). Data were analyzed by three-way ANOVA (Age × Sex × Genotype) followed by Bonferroni’s multiple-comparisons test.. *p < 0.05, **p < 0.01, ****p < 0.0001.For IL-3R⁺ microglia, significant main effects of age, sex, and genotype were detected (all p < 0.0001), along with significant Age × Genotype (p < 0.0001), Sex × Genotype (p = 0.0095), and Age × Sex × Genotype (p < 0.0001) interactions; the Age × Sex interaction alone was not significant (p = 0.0819). For IL-3⁺ astrocytes, significant main effects of age and genotype (both p < 0.0001) were observed, together with significant Age × Sex, Age × Genotype, Sex × Genotype, and Age × Sex × Genotype interactions, indicating age-dependent, genotype-specific, and sex-modulated regulation of astrocytic IL-3 expression. For GFAP⁺ astrocytes, significant main effects of age (p < 0.0001) and genotype (p = 0.0004) were detected, whereas sex alone was not significant (p = 0.8618). Significant Age × Sex, Age × Genotype, Sex × Genotype, and Age × Sex × Genotype interactions (all p ≤ 0.0002) indicate that astrocyte GFAP expression is differentially modulated by aging in a sex- and genotype-dependent manner.

To determine whether elevated microglial IL-3R expression was accompanied by parallel regulation of astrocytic cytokine production, IL-3 expression in astrocytes was assessed (**Figure 4C, D**). In female HLA mice, IL-3⁺ astrocytes increased significantly with age, rising from ∼0.5% at 6 months to ∼1% at 9 months and reaching ∼4% at 15 months, a response significantly greater than that observed in WT females or HLA males. Male HLA mice exhibited a more modest increase, from ∼1% at 6 months to ∼2% at 15 months. In contrast, WT mice of both sexes showed no age-dependent change in IL-3⁺ astrocyte frequency. These findings indicate selective, age-dependent amplification of astrocytic IL-3 expression in HLA-DRB1*15:01 mice, with a significant strongest effect in females (Figure 4d).

Astrocyte reactivity was assessed by quantifying GFAP⁺ astrocytes (**Figure 4E, F**). In WT females, GFAP⁺ astrocyte frequencies were highest at 6 and 9 months (∼30–35%) and declined significantly by 15 months (∼18%). In contrast, female HLA mice did not exhibit a comparable age-associated decline and showed significantly lower GFAP⁺ astrocyte frequencies than age-matched WT females at 6 and 9 months. In males, GFAP⁺ astrocyte frequencies remained relatively stable across age and genotype (∼20–25%). As a result, marked sex differences were evident in WT mice at younger ages (6 and 9 months), with females exhibiting higher GFAP⁺ astrocyte levels than males, whereas in HLA mice, females exhibited lower GFAP⁺ astrocyte levels than males at older ages (9 and 15 months) (**Figure 4E, F**). These patterns indicate genotype-dependent modulation of astrocyte reactivity across the lifespan.

To determine whether the IL-3R axis and activation-associated microglial markers identified by flow cytometry were also evident within intact tissue at advanced age, we performed hippocampal immunofluorescence analyses in 15-month-old WT and HLA-DRB1*15:01 mice (**Figure 5**). In WT females and males, IBA1⁺ microglia were sparsely distributed and exhibited a predominantly ramified morphology, characterized by small cell bodies with long, thin, highly branched processes consistent with a homeostatic, surveying microglial state. In these WT mice, IL-3R and CD68 immunoreactivities were minimal. In contrast, HLA mice of both sexes exhibited a marked loss of microglial ramification, with cells displaying a less branched, more amoeboid morphology indicative of a reactive, non-homeostatic state. This morphological shift was accompanied by increased IL-3R and CD68 immunoreactivity within hippocampal fields, with the strongest signal observed in HLA females and an intermediate pattern in HLA males. Merged images showed that IL-3R and CD68 signals were frequently detected within the same IBA1⁺ microglial profiles in HLA mice, whereas overlap among these markers was limited in WT mice. Together, these findings provide tissue-level confirmation that elevated microglial IL-3R expression and CD68 labeling co-occur in situ at 15 months, consistent with the age- and sex-biased patterns detected by flow cytometry.

**Figure 5.**
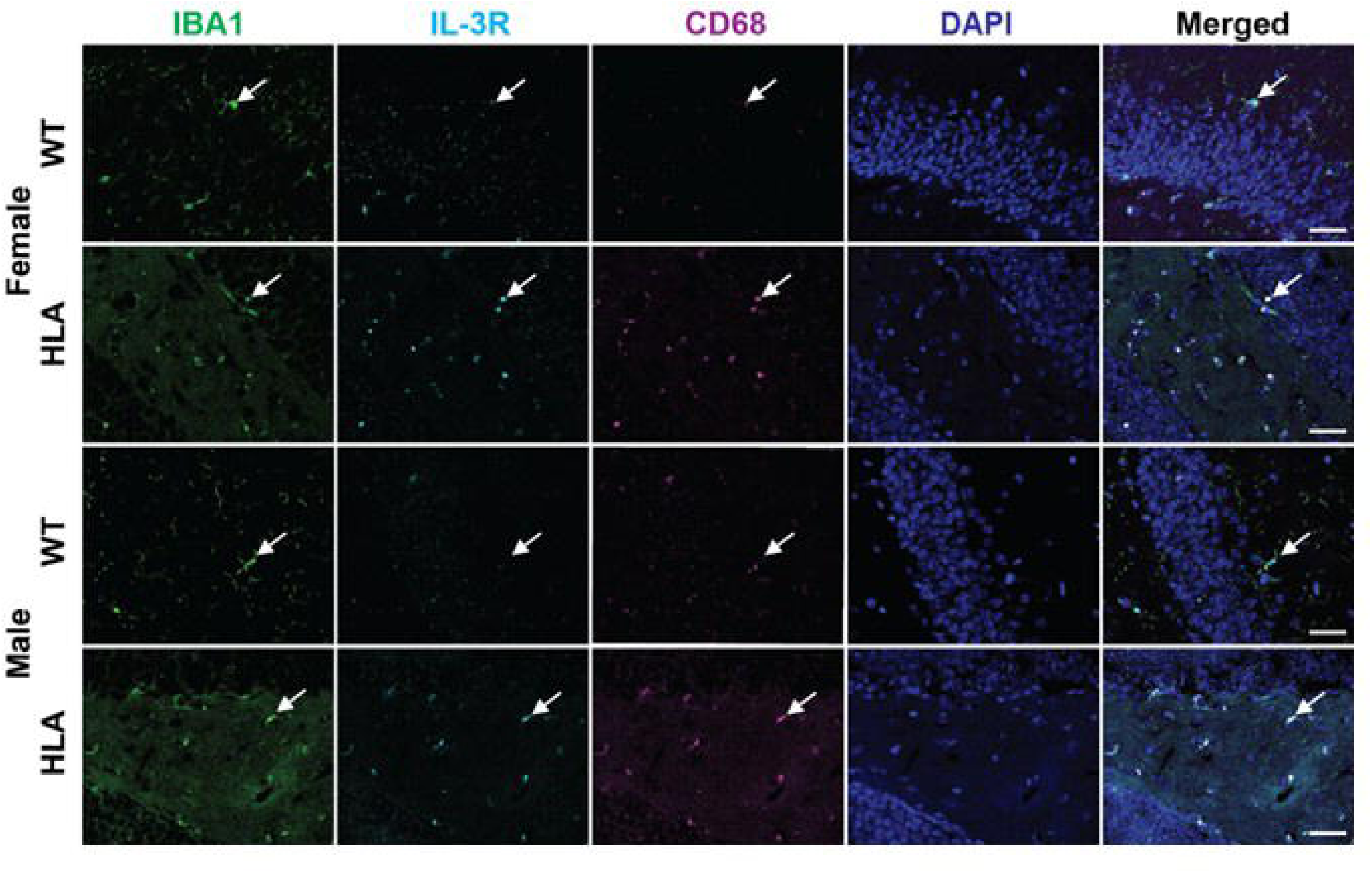
Hippocampal microglial IL-3R and CD68 immunofluorescence in aged wild-type and HLA-DRB1*15:01 mice. Representative immunofluorescence images of hippocampal sections from 15-month-old female and male wild-type (WT) and HLA-DRB1*15:01 (HLA) mice. Sections were stained for IBA1 (microglia, green), IL-3 receptor (IL-3R, cyan), and the lysosomal/phagolysosomal marker CD68 (magenta), with nuclei counterstained with DAPI (blue). Rows correspond to female WT, female HLA, male WT, and male HLA mice. White arrows indicate representative microglial cells. Images are shown as single-channel and merged views to illustrate cellular localization and marker distribution within the hippocampus. Scale bars: 20 μm.

Together, these findings demonstrate that HLA-DRB115:01 is associated with age- and sex-dependent alterations in astrocyte–microglia immune signaling. These changes include coordinated increases in IL-3–producing astrocytes and IL-3R–expressing microglia, with the largest effects observed in aged females, alongside microglial features consistent with an activated state, such as reduced ramification and increased CD68 expression. Notably, these signaling and microglial changes occurred without a parallel increase in overall astrocyte reactivity, as GFAP⁺ astrocyte frequencies were not uniformly elevated. Collectively, these results indicate selective remodeling of astrocyte–microglia immune communication in HLA-DRB115:01 mice that emerges with aging and differs by sex.

### 3.5 HLA-DRB1*15:01 expression is associated with age- and sex-dependent alterations in hippocampal microglial, neuronal, and myelin organization

Immune signaling in the brain unfolds within a structured cellular environment in which the spatial relationships among glial cells, neurons, and myelin shape neuroimmune interactions. Sustained or dysregulated immune signaling can remodel tissue architecture, and we therefore examined whether the age- and genotype-dependent microglial and astrocytic phenotypes identified earlier were accompanied by corresponding changes in hippocampal cellular organization. Using immunofluorescence in 15-month-old WT and HLA mice, we assessed the spatial relationships among microglia, axonal structure, myelin, and astrocytic markers (GFAP and IL-3) within the hippocampus.

As shown in Figure 6, and consistent with Figure 5, WT mice of both sexes displayed sparsely distributed microglia with a predominantly ramified morphology (IBA1⁺), whereas HLA mice exhibited less ramified microglial profiles. NF200 immunostaining, used to visualize axonal structure, revealed qualitative genotype-dependent differences. WT females and males showed dense, continuous, and well-organized axonal labeling. In contrast, HLA females displayed reduced and discontinuous NF200 signal, suggesting more fragmented axonal profiles. HLA males exhibited a comparable but less pronounced reduction relative to WT males, indicating a sex-dependent effect (**Figure 6A**).

**Figure 6.**
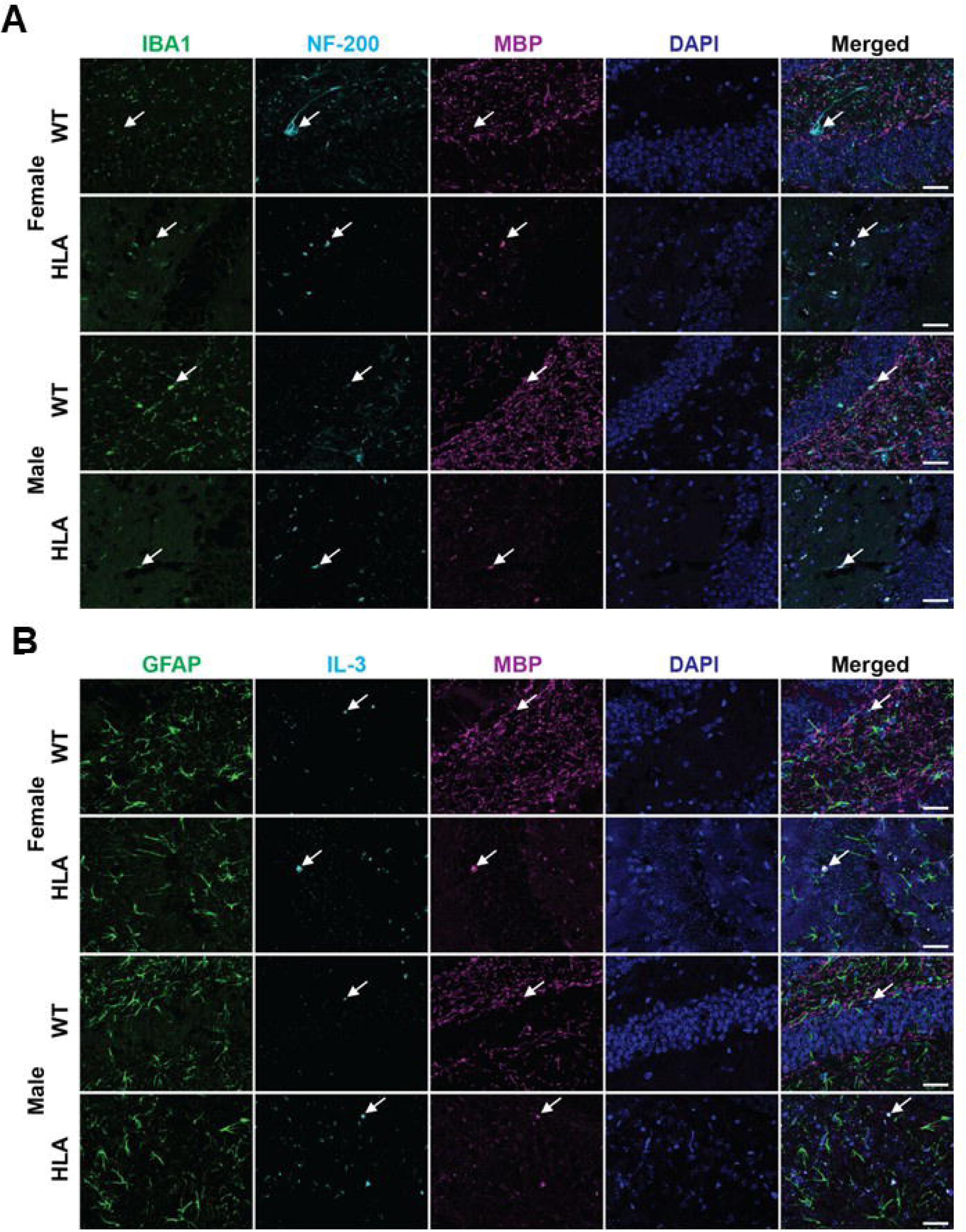
Tissue-level organization of microglia, axonal structure, myelin, and astrocytic IL-3 signaling in the aged hippocampus of wild-type and HLA-DRB1*15:01 mice. Hippocampal sections from 15-month-old female and male wild-type (WT) and HLA-DRB1*15:01 (HLA). (A) Representative immunofluorescence images showing microglia (IBA1, green), axonal structure labeled by NF200 (cyan), myelin (MBP, magenta), and nuclei (DAPI, blue) in the hippocampus. Merged images illustrate the spatial relationship between microglia, axonal profiles, and myelinated structures. White arrows indicate representative microglial cells, axonal, and myelin structure, as appropriate. (B) Representative immunofluorescence images from the same hippocampal regions showing astrocytes labeled by GFAP (green), IL-3 immunoreactivity (cyan), myelin (MBP, magenta), and nuclei (DAPI, blue). White arrows indicate representative GFAP⁺ astrocytes or IL-3⁺ puncta, as appropriate. Images are shown as single-channel and merged views to illustrate cellular localization and marker distribution within the hippocampus. Scale bars, 20 µm.

Myelin immunostaining (MBP) also revealed genotype-associated differences in hippocampal organization. WT females and males displayed strong, continuous myelin labeling with clearly defined fiber-like structures. In contrast, both HLA females and HLA males exhibited reduced and more diffuse myelin immunoreactivity, characterized by diminished organization and weaker overall signal in representative fields (**Figure 6A, B**). Merged images integrating microglia, axonal, and myelin channels showed that regions of reduced axonal and myelin immunoreactivity in HLA mice frequently coincided with areas of increased microglial presence. This spatial relationship was more prominent in HLA females than in HLA males. In WT mice of both sexes, neuronal and myelin architecture appeared preserved, with microglia largely interspersed among intact tissue structures without showing co-localization (**Figure 6A**).

GFAP immunoreactivity in WT and HLA mice of both sexes showed dense, widespread labeling across the hippocampal field. IL-3 immunoreactivity displayed a distinct genotype-dependent pattern. HLA females and males exhibited increased IL-3 signal, with a greater number of discrete IL-3⁺ puncta distributed throughout the tissue compared with WT mice. Across all groups, merged images revealed no overlap between GFAP⁺ astrocytes and IL-3 immunoreactivity. Notably, in HLA mice of both sexes, IL-3 signal was frequently observed within regions exhibiting reduced myelin organization, a co-localization not evident in WT mice (**Figure 6B**).

Together, these findings indicate that HLA-DRB115:01 expression is associated with coordinated, age-and sex-dependent alterations in hippocampal cellular organization at 15 months of age. In HLA mice, reduced microglial ramification and increased microglial presence spatially coincide with disrupted axonal and myelin organization, with these changes more pronounced in females. While overall astrocytic GFAP labeling appeared preserved across genotypes, HLA mice exhibited a distinct increase in IL-3 immunoreactivity that localized preferentially to regions with reduced myelin structure and showed no overlap with GFAP⁺ astrocytes. These observations define a genotype-dependent spatial remodeling of immune-associated signals within the hippocampus, providing tissue-level context for the cellular and molecular neuroimmune alterations observed in HLA-DRB115:01 mice.

### 3.6 HLA-DRB1*15:01 drives age- and sex-dependent immune cell recruitment and endothelial activation at CNS borders

Neuroinflammatory processes in the aging brain are shaped not only by resident glial responses but also by tightly regulated immune cell trafficking across CNS interfaces and by endothelial activation that governs leukocyte entry. Recent work has identified astrocyte-derived interleukin-3 (IL-3) as a key amplifier of neuroimmune crosstalk. Early IL-3 production by activated astrocytes can stimulate IL-3 receptor–expressing microglia to produce chemokines that promote the recruitment of CD4⁺ T cells into the CNS. Given the age- and sex-dependent alterations in microglial activation and astrocyte–microglia IL-3/IL-3R signaling observed in HLA-DRB1*15:01 mice, we next examined whether these changes were accompanied by selective lymphoid immune cell accumulation at CNS borders and within the brain parenchyma, as well as by endothelial activation at the neurovascular interface. To address this, we quantified CD3⁺ T cells, CD4⁺ T-cell subsets, and NK1.1⁺CD3⁺ natural killer T (NKT) cells in the meninges and brain across age, sex, and genotype, and assessed endothelial adhesion molecule expression associated with leukocyte recruitment. CD8⁺ T cells were not detected at appreciable levels in either compartment and were therefore excluded from further analysis.

We quantified the percentage of total T cells in the meninges and brain across age, sex, and genotype (**Figure 7A, B**). In female mice, 15-month-old HLA animals showed a significant increase in meningeal T cell frequency (∼6.5%) compared with 6-month-old HLA females (∼1.5%; p < 0.0001), 9-month-old HLA females (∼3.5%; p = 0.0004), and age-matched WT females (WT ∼2% vs HLA ∼6.5%; p < 0.0001) (**Figure 7A**). Comparison between sexes revealed a significant sex-dependent effect at 15 months, with HLA females exhibiting higher meningeal T cell frequencies than HLA males (female ∼6.5% vs male ∼3.5%; p = 0.004). No significant differences were detected among the remaining age, sex, or genotype groups in male and female WT mice or in male HLA mice (**Figure 7A**). In contrast, analysis of brain-infiltrating T cells revealed no statistically significant differences across age, sex, or genotype after correction for multiple comparisons (**Figure 7B**). Exploratory post hoc (Fisher’s LSD) analyses identified pairwise differences in brain T cell percentages in 15-month-old HLA mice of both sexes compared with WT controls (female: HLA ∼1.7% vs WT ∼0.9%, p = 0.0087; male: HLA ∼1.8% vs WT ∼1.2%, p = 0.02); however, these differences did not remain significant following correction (Bonferroni/Sidak) and should therefore be interpreted as exploratory.

**Figure 7.**
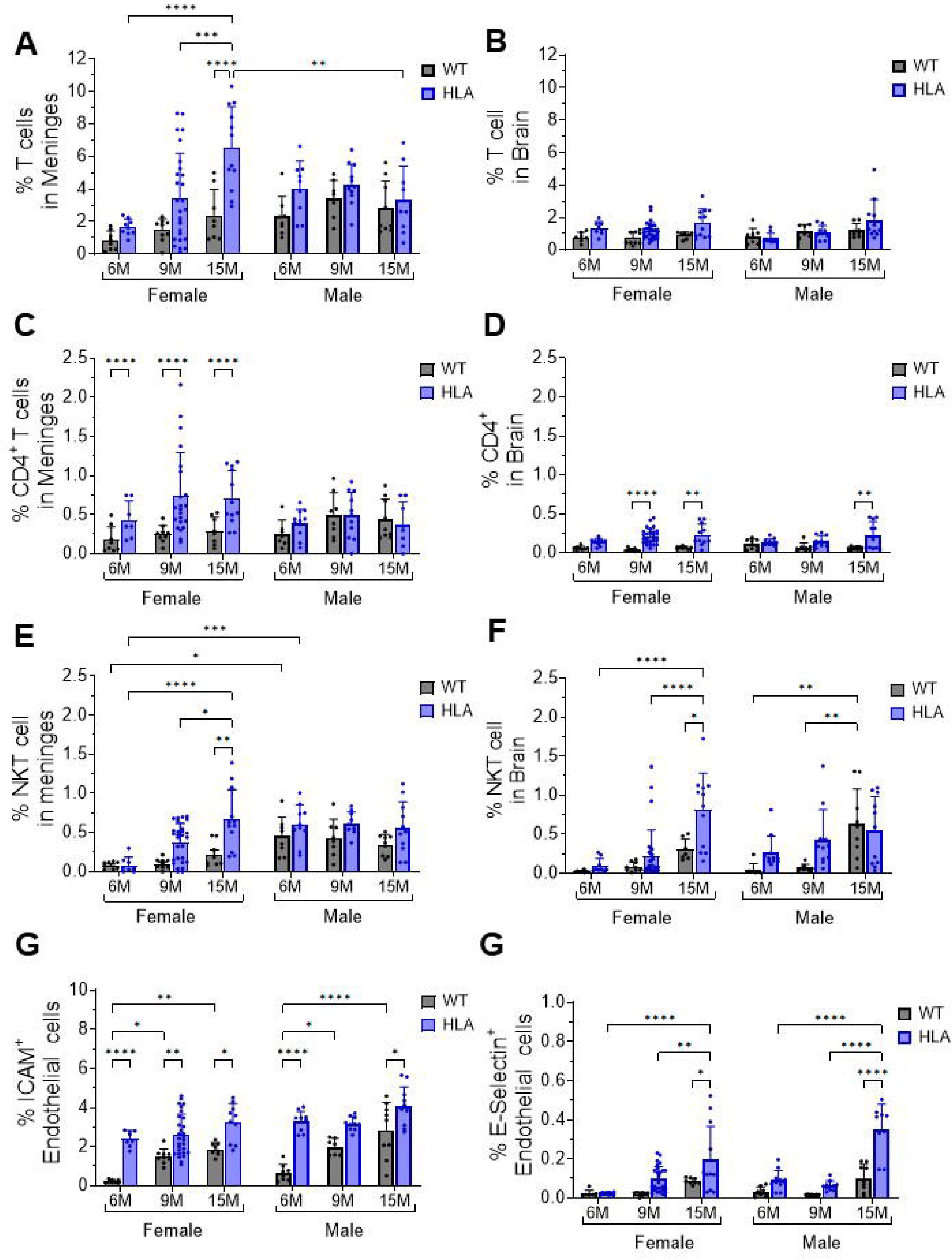
Age- and sex-dependent T cell accumulation and endothelial activation in the meninges and brain of HLA-DRB1*15:01 mice. Meninges and brain tissue were dissociated from female and male wild-type (WT, gray) and HLA-DRB1*15:01 (HLA, blue) mice at 6, 9, and 15 months of age, stained with cell type–specific markers, and analyzed by flow cytometry. Cells were gated based on forward- and side-scatter characteristics, with dead cells and doublets excluded using DAPI staining and FSC-A/FSC-H parameters. T cells were identified as CD45⁺CD3⁺ cells, and CD4⁺ T cells were defined as CD45⁺CD3⁺CD4⁺CD8⁻ cells. (A) Percentage of total T cells and (B) percentage of CD4⁺ T cells in the meninges. (C) Percentage of total T cells and (D) percentage of CD4⁺ T cells in the brain. Endothelial cells were identified as CD31⁺ cells, and endothelial activation was assessed by adhesion molecule expression. (E) Percentage of E-selectin⁺ (CD62E⁺) endothelial cells and (F) percentage of ICAM-1⁺ (CD54⁺) endothelial cells in the brain. Data are presented as mean ± SEM with individual mice shown. Statistical significance was determined by three-way ANOVA followed by Bonferroni’s multiple-comparisons test (*p < 0.05, **p < 0.01, ***p < 0.001, ****p < 0.0001). For meningeal CD3⁺ T cells, ANOVA revealed significant main effects of age (p = 0.0027) and genotype (p < 0.0001), whereas sex alone was not significant. Significant Age × Sex (p = 0.0004) and Age × Sex × Genotype (p = 0.0380) interactions were detected, while Age × Genotype and Sex × Genotype interactions were not significant. For meningeal CD3⁺CD4⁺ T cells, a significant main effect of genotype (p = 0.0012) and a significant Sex × Genotype interaction (p = 0.0035) were observed, with no significant effects of age or higher-order interactions. For brain CD4⁺ T cells, a significant main effect of genotype was detected (p < 0.0001), whereas age and the Age × Genotype interaction were not significant. For E-selectin⁺ endothelial cells in the brain, three-way ANOVA revealed significant main effects of age, sex, and genotype (all p < 0.05), along with significant Age × Sex, Age × Genotype, Sex × Genotype, and Age × Sex × Genotype interactions. For ICAM-1⁺ endothelial cells, significant main effects of age, sex, and genotype were observed (all p < 0.0001), together with a significant Age × Genotype interaction (p = 0.0017); no other interactions reached significance.

Female HLA mice also exhibited a significant increase in the percentage of meningeal CD4⁺ T cells compared with age-matched WT controls at 6, 9, and 15 months (all p < 0.0001) (**Figure 7C**). However, no age-dependent increase was observed within the HLA female group. In contrast, neither WT nor HLA male mice showed significant differences in meningeal CD4⁺ T cell frequencies across age or genotype (**Figure 7C**). Analysis of CD4⁺ T cells in the brain also revealed significant genotype-dependent differences (**Figure 7D**). Female HLA animals showed a significant increase in brain CD4⁺ T cell frequency at 9 and 15 months compared with age-matched WT controls (p < 0.01 and p < 0.0001, respectively) (**Figure 7D**). HLA males showed a significant increase in brain CD4⁺ T cell frequency at 15 months relative to WT males (p < 0.01), whereas no significant differences were detected at earlier ages (**Figure 7D**).

Analysis of NK1.1⁺CD3⁺ natural killer T (NKT) cells in the meninges revealed significant age-, sex-, and genotype-dependent differences (**Figure 7E**). Compared with age-matched WT controls, HLA females exhibited significantly higher percentages of meningeal NKT cells at 15 months (p = 0.014). Within the HLA female group, NKT cell levels at 15 months were significantly elevated relative to both 6 months (p < 0.0001) and 9 months (p = 0.012) (**Figure 7E**). In contrast, HLA male mice did not exhibit significant age- or genotype-dependent differences in meningeal NKT cell frequencies. A significant sex difference was observed at 6 months in both genotypes, with males exhibiting higher percentages of meningeal NKT cells than females in both the WT (p = 0.04) and HLA (p = 0.0002) groups (**Figure 7E**). In the brain parenchyma, NKT cell analysis did not reveal a uniform sex-independent effect across genotypes (**Figure 7F**). However, similar to the meninges, brain NKT cells showed significant age- and genotype-dependent effects in females but not in males. In HLA females, NKT cell levels at 15 months were significantly higher than at both 6 months (p < 0.0001) and 9 months (p < 0.0001), and were also elevated relative to age-matched WT controls (p = 0.016) (Figure 7f). In contrast, HLA males did not exhibit a significant age-dependent effect. Notably, WT males showed increased brain NKT cell levels at 15 months compared with both 6 months (p = 0.002) and 9 months (p = 0.0052) (**Figure 7F**).

Endothelial activation in the brain was assessed by analyzing the expression of adhesion molecules, including P-selectin, E-selectin, VCAM-1, and ICAM-1. P-selectin and VCAM-1 were not detected at appreciable levels; therefore, subsequent analyses focused on ICAM-1 and E-selectin as indicators of endothelial activation at the neurovascular interface (**Figure 7G, H**). Analysis of ICAM-1–positive endothelial cells revealed significant age- and genotype-dependent effects (**Figure 7G**). HLA females and males did not exhibit significant age- or sex-dependent differences in ICAM-1⁺ endothelial cell percentages within the HLA genotype. However, ICAM-1 expression was consistently higher in HLA mice compared with age-matched WT controls in both sexes. In females, ICAM-1⁺ endothelial cells were significantly increased in HLA mice at 6 months (∼2.5% vs ∼0.3%; p < 0.0001), 9 months (∼2.5% vs ∼1.5%; p = 0.008), and 15 months (∼3.0% vs ∼2.0%; p = 0.05). Similarly, in males, HLA mice showed significantly higher ICAM-1 expression at 6 months (∼3.0% vs ∼0.5%; p < 0.0001) and 15 months (∼4.0% vs ∼3.0%; p < 0.0001) compared with WT controls. In contrast, WT females and males showed an age-associated increase in ICAM-1 expression, with significantly higher levels at 15 months than at 6 months in both sexes. E-selectin–positive endothelial cells were detected at much lower frequencies than ICAM-1–positive cells (**Figure 7G, H**). Nevertheless, E-selectin analysis revealed significant age- and genotype-dependent differences (**Figure 7H**). In female mice, HLA animals exhibited a significant increase in E-selectin⁺ endothelial cells at 15 months compared with 6 months (p < 0.0001), 9 months (p = 0.0059), and age-matched WT females (p < 0.001). In male mice, E-selectin⁺ endothelial cells were similarly increased in 15-month-old HLA animals compared with 6 months (p < 0.0001), 9 months (p < 0.0001), and age-matched WT males (p < 0.0001). No significant differences were detected at earlier ages.

Together, these findings demonstrate that HLA-DRB1*15:01 expression promotes selective, age- and sex-dependent immune remodeling at CNS interfaces, rather than generalized immune infiltration. Increased accumulation of CD4⁺ T cells and NKT cells—particularly in females—occurs alongside endothelial activation characterized by elevated ICAM-1 and E-selectin expression, suggesting a permissive neurovascular environment for regulated immune cell recruitment. When considered in the context of the preceding results showing microglial activation, IL-3/IL-3R signaling, and tissue-level remodeling, these data support a model in which HLA-DRB115:01 amplifies chronic, compartmentalized neuroinflammatory processes that emerge with aging and are strongly modulated by sex.

### 3.7 HLA-DRB1*15:01 expression alters selective brain cytokine signaling with aging

Neuroinflammatory outcomes are shaped not only by cellular composition and immune trafficking but also by the local cytokine environment within the brain parenchyma. Given the age- and sex-dependent alterations in glial activation, IL-3/IL-3R signaling, endothelial activation, and T cell accumulation observed in HLA-DRB1*15:01 mice, we next examined whether these cellular changes were accompanied by corresponding shifts in brain cytokine profiles. To address this, cytokine levels were quantified in lysed hippocampal tissue from WT and HLA mice across age and sex using a multiplex electrochemiluminescence assay (MSD). The cytokine panel included IFN-γ, IL-1β, IL-2, IL-4, IL-5, IL-6, IL-10, IL-12p70, KC/GRO, and TNF-α. Among the cytokines assessed, only IL-12p70, IL-10, and IL-2 exhibited detectable and statistically analyzable differences across age, sex, or genotype. Levels of IFN-γ, IL-1β, IL-4, IL-5, IL-6, KC/GRO, and TNF-α did not differ significantly between groups or were below the assay’s reliable detection range. Accordingly, subsequent analyses focused on IL-12p70, IL-10, and IL-2 (**Figure 8**).

**Figure 8.**
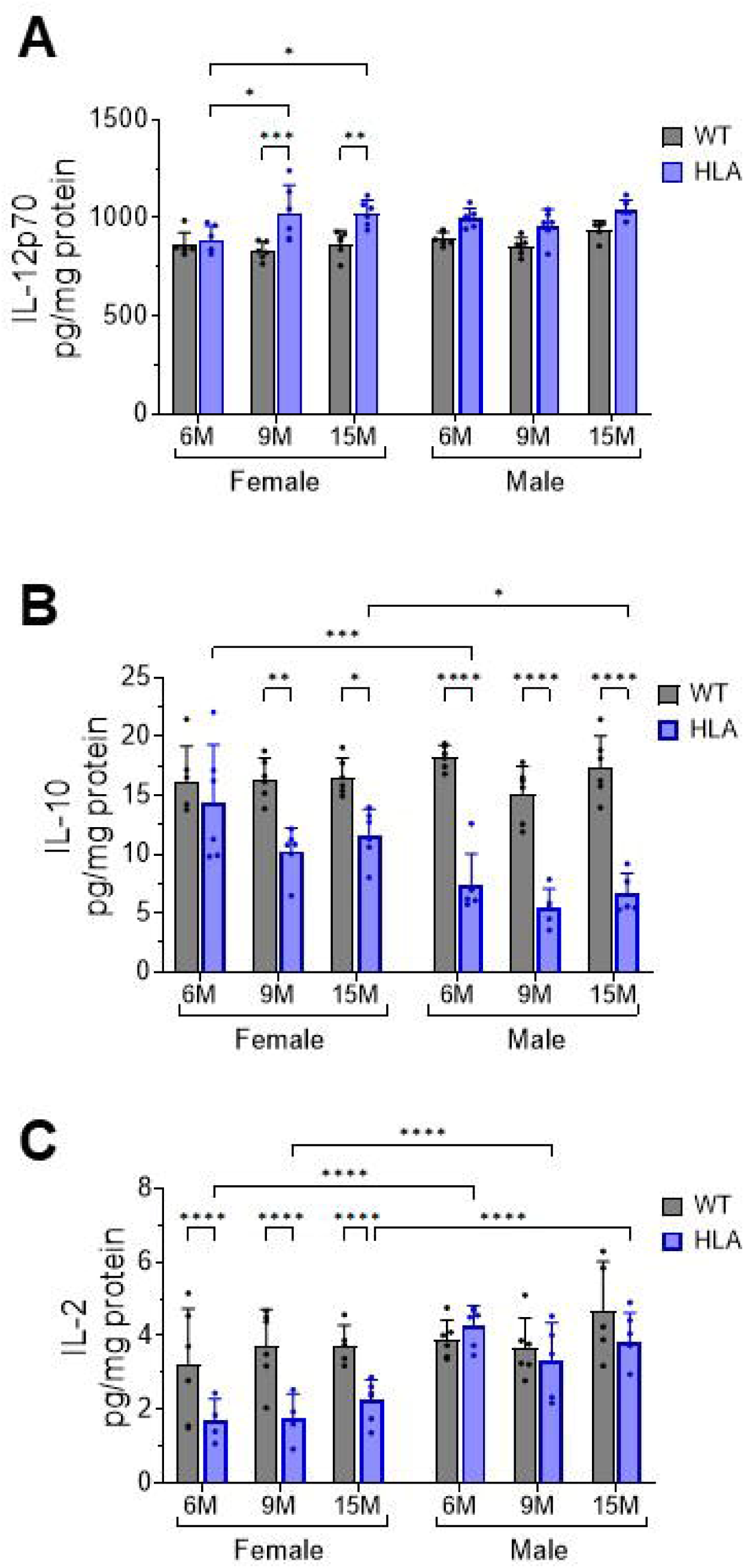
Selective alterations in hippocampal cytokine profiles with aging in HLA-DRB1*15:01 mice. Hippocampal tissue was collected from female and male wild-type (WT, gray) and HLA-DRB1*15:01 (HLA, blue) mice at 6, 9, and 15 months of age. Tissue was lysed, and cytokine concentrations were quantified using a multiplex electrochemiluminescence assay (MSD). Among the cytokines assessed, only IL-12p70, IL-10, and IL-2 were detected at levels suitable for statistical analysis. (A) IL-12p70, (B) IL-10, and (C) IL-2 concentrations are shown and expressed as pg/mg total protein. Data are presented as mean ± SEM with individual mice shown. Statistical significance was determined by three-way ANOVA followed by Bonferroni’s multiple-comparisons test. *p < 0.05, **p < 0.01, ***p < 0.001, ****p < 0.0001. IL-12p70 showed a significant main effect of age (p = 0.0109) and a significant main effect of genotype (p < 0.0001). IL-10 revealed significant main effects of age (p = 0.0103), sex (p < 0.0001), genotype (p < 0.0001), and a significant Sex × Genotype (p < 0.0001). IL-2 showed significant main effects of sex and genotype (both p < 0.0001) and a significant Sex × Genotype interaction (p = 0.0031).

Analysis of IL-12p70 levels in hippocampal lysates revealed age- and genotype-dependent differences specifically in female mice. In females, HLA mice exhibited significantly higher IL-12p70 levels at 9 months (p = 0.0002) and 15 months (p = 0.0065) compared with age-matched WT controls (Figure 8a). In addition, IL-12p70 levels in HLA females increased with age, with significantly higher levels at 9 and 15 months compared with 6 months (p = 0.02). In contrast, male mice showed no significant genotype- or age-dependent differences in IL-12p70 levels across the time points examined (**Figure 8A**).

Quantification of IL-10 levels revealed a strong genotype-dependent reduction in HLA mice, with clear sex-specific differences. In female mice, IL-10 levels were significantly lower in HLA animals than in age-matched WT controls at 9 months (p = 0.001) and 15 months (p = 0.025), whereas no significant difference was detected at 6 months (**Figure 8B**). In male mice, IL-10 levels were significantly lower in HLA animals than in WT controls at all ages examined (6, 9, and 15 months; all p < 0.0001) (**Figure 8B**). IL-10 levels did not exhibit significant age-dependent changes within either WT or HLA groups in females or males. Comparison across sexes further indicated that IL-10 levels were significantly lower in HLA males than in HLA females (**Figure 8B**).

Quantification of IL-2 levels in hippocampal lysates revealed significant genotype- and sex-dependent differences within the HLA genotype (**Figure 8C**). In female mice, IL-2 levels were consistently lower in HLA-DRB1*15:01 animals compared with age-matched WT controls at all ages examined (all p < 0.0001). In contrast, no significant genotype-dependent differences were detected in male mice. IL-2 levels did not show significant age-associated changes within either genotype in males or females. Comparison across sexes revealed a pronounced sex difference within the HLA genotype, with HLA males exhibiting significantly higher IL-2 levels than HLA females across ages (**Figure 8C**).

Together, these results indicate that HLA-DRB1*15:01 expression selectively alters the hippocampal cytokine environment in an age- and sex-dependent manner. Increased IL-12p70 in aged HLA females, combined with reduced IL-10 and suppressed IL-2 in HLA females, reflects a shift toward pro-inflammatory and dysregulated immune signaling within the aging brain. When considered alongside earlier findings of glial activation, IL-3/IL-3R signaling, endothelial activation, and increased CD4⁺ T-cell accumulation, these cytokine changes support a coordinated neuroinflammatory phenotype associated with HLA-DRB1*15:01 during aging.

## 4 DISCUSSION

The present study demonstrates that expression of the human autoimmune risk allele HLA-DRB1*15:01—a genetic risk factor for MS and susceptibility to late-onset neurodegenerative diseases —is associated with selective, age- and sex-dependent remodeling of the neuroimmune environment in the aging brain. By integrating behavioral analyses with flow cytometry, immunofluorescence, and hippocampal cytokine profiling, we identify a coordinated HLA-DRB1*15:01–associated phenotype characterized by microglial oxidative stress and activation-related changes, altered astrocyte–microglia immune signaling, endothelial activation at CNS interfaces, selective lymphoid cell accumulation, and targeted cytokine imbalance. Notably, these effects were most pronounced in aged females and coincided with the emergence of late-onset behavioral impairment, underscoring a sex-selective neuroimmune vulnerability associated with HLA-DRB1*15:01.

It is important to point out that in this study, HLA-DRB115:01 mice were not challenged with myelin peptides such as mMOG-35–55 and did not develop clinical paralysis or overt autoimmune demyelination. In addition, this humanized model lacks endogenous murine MHC class II, restricting classical murine MHC-II–dependent antigen presentation by CNS-resident and peripheral antigen-presenting cells (22). Consequently, the observed phenotypes reflect baseline, age-related neuroimmune changes associated with HLA-DRB1*15:01, rather than effects of active autoimmune disease. At the same time, this represents an important caveat: the immune architecture of this model is not identical to that of WT mice with intact murine MHC-II, and thus the phenotype should be viewed as HLA-driven remodeling within an MHC-II–null context.

A key finding of this study is the emergence of late-onset cognitive impairment restricted to aged female HLA-DRB1*15:01 mice. At 15 months of age, HLA females, but not males, exhibited deficits in nest building, recognition memory, and spatial discrimination, while WT mice of both sexes remained behaviorally intact. The delayed onset of impairment argues against early neurodevelopmental dysfunction and instead supports a model in which HLA-DRB1*15:01 interacts with aging-related processes, particularly in females. This sex bias aligns directionally with immune-mediated CNS disorders such as MS, in which sex and age shape susceptibility and disease trajectories (35, 36). Importantly, behavioral impairment temporally coincided with the strongest convergence of neuroimmune alterations, supporting the interpretation that functional decline arises within a progressively remodeled immune environment rather than from a single acute inflammatory trigger.

Microglial oxidative stress emerged as an early and robust feature of HLA-DRB1*15:01 expression. Both female and male HLA mice displayed age-dependent increases in MitoSOX-positive microglia beginning at midlife, whereas WT mice maintained low oxidative burden across ages. At advanced age, HLA females exhibited the highest levels of oxidative stress. Oxidative stress, defined as an imbalance between the production and clearance of reactive oxygen species, is closely linked to metabolic stress in microglia and can influence cellular homeostasis and immune responsiveness (37, 38). While oxidative stress alone cannot be interpreted as definitive evidence of microglial priming, accumulating evidence indicates that an altered redox state can modulate microglial functional properties and responsiveness to subsequent immune cues (39–41). In this context, the emergence of elevated oxidative stress at 9 months, preceding behavioral impairment at 15 months in females, suggests that redox imbalance represents an early alteration in microglial state, rather than a downstream consequence of behavioral decline. . Because microglial oxidative stress emerges at midlife, ahead of behavioral impairment and immune cell accumulation, it may reflect an early permissive condition that influences later neuroimmune remodeling.

Consistent with this oxidative phenotype, HLA-DRB1*15:01 expression was associated with progressive, sex and age-dependent increases in activation-associated microglial markers measured using MHC-II independent readouts (CD14 and CD68). This strategy was necessary given the absence of endogenous murine MHC class II and enabled genotype comparisons without relying on MHC-II expression. CD14 upregulation is consistent with heightened innate immune responsiveness (32, 33), while increased CD68 reflects enhanced lysosomal and phagolysosomal activity characteristic of activated microglia (34). These changes were not uniform across age or sex, but instead microglial activation was most pronounced in aged females. At the tissue level, immunofluorescence revealed reduced microglial ramification. Under homeostatic conditions, ramified microglia exhibit highly dynamic process motility and continuously survey neuronal somata and synapses, a defining feature of physiological microglial function (42–44). In contrast, chronic neurodegenerative and inflammatory states are associated with a transition to a reactive phenotype characterized by loss of ramified architecture, reflecting altered surveillance and engagement with the neural environment (42–44). Although morphology alone does not define a specific functional program, the concurrent loss of ramification, increased CD68 expression, and elevated oxidative stress observed in HLA mice is consistent with a shift away from a homeostatic surveying state toward a sustained reactive microglial phenotype. Taken together, these findings indicate that HLA-DRB1*15:01 biases microglia toward a metabolically stressed, reactive state that emerges gradually with aging.

A mechanistically-related aspect of this study is the coordinated, age- and sex-dependent amplification of the IL-3/IL-3 receptor axis across astrocytes and microglia. IL-3R expression on microglia increased with age in both genotypes but was markedly elevated in HLA-DRB1*15:01 mice, with the strongest induction observed in aged females. Notably, MS exhibits a strong female bias, and the IL3RA gene is encoded on the X chromosome (45), raising the possibility that sex-linked regulatory mechanisms may contribute to the enhanced IL-3R expression observed in HLA females. In parallel, astrocytic IL-3 expression increased selectively in HLA mice, again with the greatest magnitude in females. This coordinated pattern is notable because accumulating evidence indicates that IL-3 functions as a local CNS immune-modulatory signal, rather than a systemic hematopoietic cytokine, shaping microglial immune tone, chemotactic programs, and regulated immune recruitment at CNS interfaces(46–53). Importantly, astrocytes have been identified as sources of IL-3 in both healthy and inflamed CNS tissue, and IL-3 expression is not linked to classical astrogliosis (48, 50). Consistent with these observations, increased IL-3 immunoreactivity in HLA mice occurred without generalized GFAP upregulation, supporting selective astrocyte functional remodeling.

The functional consequences of IL-3 signaling are highly context dependent. In Alzheimer’s disease, IL-3 promotes microglial clustering and clearance of amyloid and tau, exerting protective effects, whereas in human MS and an EAE murine model, IL-3 signaling exacerbates disease by promoting immune recruitment and neuroinflammation (50, 54). These contrasting outcomes underscore that the impact of IL-3/IL-3R signaling depends on cellular targets, disease environment, and immune architecture. In the present study, IL-3/IL-3R amplification emerged in the absence of EAE induction, supporting the concept that genetic risk conferred by HLA-DRB1*15:01 may modulate IL-3–mediated signaling as part of baseline, age-associated immune tuning within the CNS, rather than as a late-stage inflammatory byproduct. Nevertheless, an important caveat is that these data are correlative. While the close coupling between astrocytic IL-3 expression, microglial IL-3R upregulation, and microglial activation strongly implicates this axis in shaping the observed phenotype, causality cannot be inferred without direct genetic or pharmacological perturbation.

Despite the absence of overt tissue pathology, HLA-DRB1*15:01 expression was associated with coordinated qualitative alterations in hippocampal organization at advanced age, including reduced axonal labeling, more diffuse myelin basic protein organization, and increased microglial presence in regions of altered structure. The more pronounced alterations observed in females align with the stronger immune phenotype in this group. However, these observations are currently qualitative and based on representative fields. Future studies incorporating quantitative morphometric analyses, systematic regional sampling, and ultrastructural approaches will be required to determine their magnitude and biological significance.

HLA-DRB1*15:01 also influenced immune dynamics at CNS borders. Rather than broad infiltration, we observed selective accumulation of CD4⁺ T cells and NKT cells in the meninges and brain, with a preference in females. Endothelial activation was also observed, characterized by increased expression of ICAM-1 and E-selectin. Together, these findings suggest a state of endothelial priming sufficient to support regulated leukocyte recruitment, without full vascular activation. In addition, the evidence that IL-3/IL-3Rα signaling can drive chemotactic programming in myeloid cells and promote CNS immune recruitment (50) raises the possibility that IL-3/IL3R signaling contributes to immune activity at CNS borders in HLA-DRB1*15:01 mice.

Finally, hippocampal cytokine profiling reinforced the theme of selectivity. Among the 10 cytokines assessed, only IL-12p70, IL-10, and IL-2 showed reliable differences across age, sex, and genotype, indicating a selective alteration of cytokine signaling rather than a global inflammatory response. The cytokine profile observed in aged female HLA mice, elevated IL-12p70 coupled with reduced IL-10 and IL-2, reflects a shift toward pro-inflammatory immune polarization with diminished regulatory balance, rather than generalized cytokine excess. Because cytokine measurements were derived from whole hippocampal lysates, cellular sources cannot be resolved, underscoring the need for future spatial and cell-specific analyses.

## 5 Conclusions

Taken together, these findings support an integrated model in which HLA-DRB1*15:01 lowers the threshold for age-related neuroimmune dysregulation rather than directly inducing autoimmune demyelination. In the absence of antigenic challenge, this allele is associated with a coordinated, sex-biased phenotype involving oxidative stress, microglial activation-associated changes, selective IL-3/IL-3R signaling, endothelial priming, compartmentalized immune recruitment, and targeted cytokine imbalance. The most compelling aspects of this work are (i) the demonstration of an age- and sex-dependent phenotype emerging under baseline conditions, and (ii) the convergence of immune remodeling across multiple levels of analysis that aligns temporally with female-restricted behavioral impairment. At the same time, interpretation must be tempered by key limitations, including the absence of murine MHC-II, the correlative nature of IL-3/IL-3R associations, and the qualitative nature of some tissue-level findings. Future studies incorporating pathway perturbation, spatial transcriptomics, and inflammatory or antigenic challenges will be essential to define how these baseline neuroimmune alterations interact with autoimmune mechanisms during disease initiation and progression.

## Author contributions

KER and EMRR contributed to experimental planning, data interpretation and analysis, and manuscript preparation. DC prepared the IHC figures and performed the analysis. EMRR, DC, FF, MDT, VDN, QH, CF, ACL, DB and JPW contributed to experimental execution.

## Ethics approval

All the animal studies were conducted in accordance with the ethical guidelines set forth by the National Institutes of Health guidelines for procedures on laboratory animals and approved by the University of Arizona Institutional Care and Use Committee.

## Generative AI statement

Generative artificial intelligence was used solely to assist with language editing.

## Acknowledgments

We thank Lisa Campbell for her contributions.

