## Supplementary Figure 1-5 for "HLA-DRB1*15:01 drives sex- and age-dependent microglial activation and neuroimmune signaling"

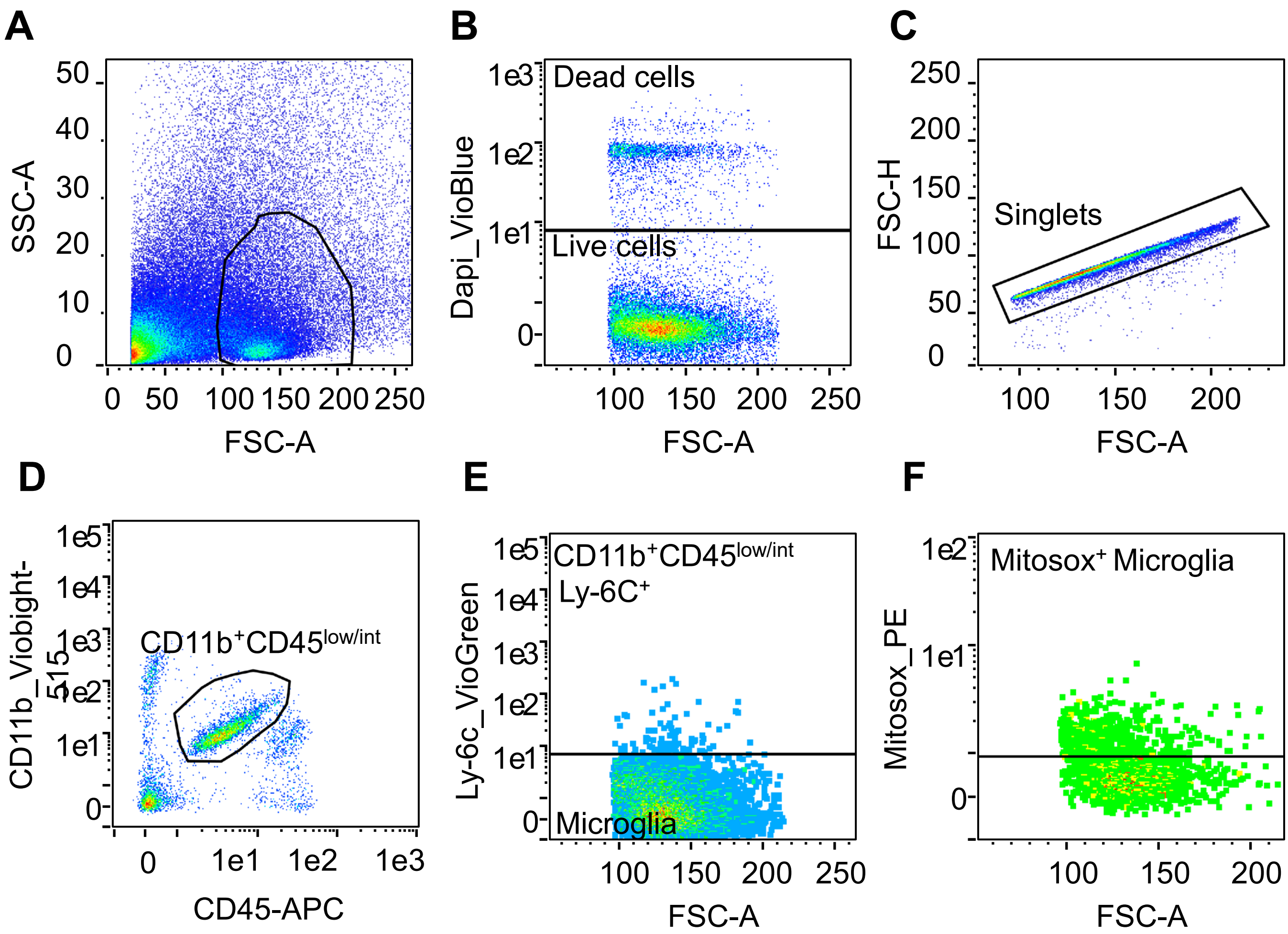


**Supplementary Figure 1. Gating strategy for identification of MitoSOX$^+$ microglia**.
(A) Single-cell suspensions from adult mouse brain were first gated on FSC-A vs SSC-A to exclude debris. (B) Dead cells were excluded using DAPI staining (C) followed by FSC-A/FSC-H gating to select singlets. (D) Microglia were identified as CD11b$^+$ CD45$^{low/int}$ cells. (E) Peripheral monocytes were further excluded based on Ly6C expression.(F) Mitochondrial superoxide production was assessed using MitoSOX-PE within the gated microglia population.

#### Supplementary Figure 2

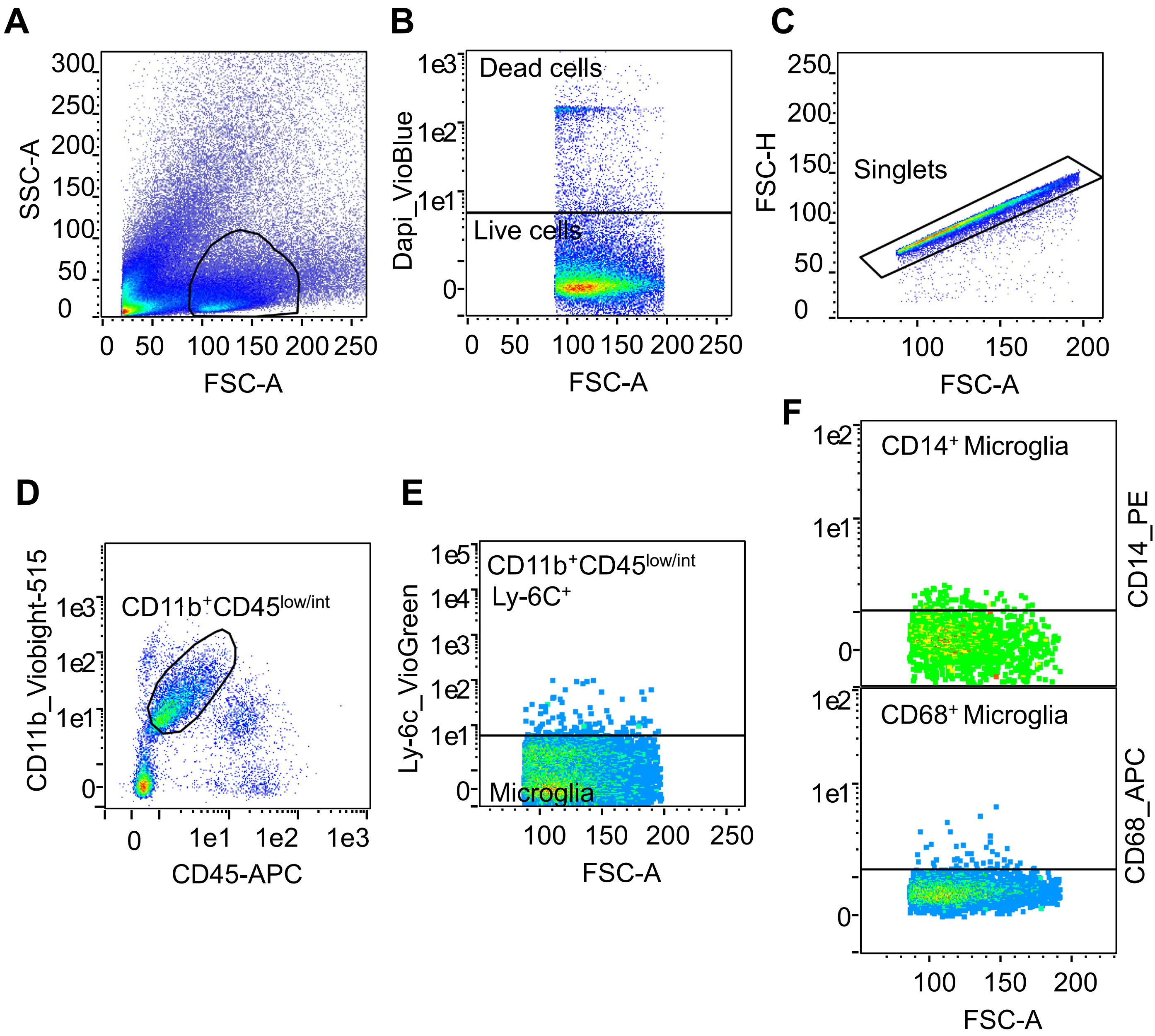


**Supplementary Figure 2. Microglia gating and activation analysis by flow cytometry.** Brain single-cell suspensions were gated to (A) exclude debris, (B) dead cells (DAPI$^+$, and (C) doublets. (D) Microglia were defined as CD11b$^+$ CD45$^{low/int}$ cells, (E) excluding Ly6C$^+$ events. Activation was assessed by (F) CD14 and CD68 expression.

### Supplementary Figure 3

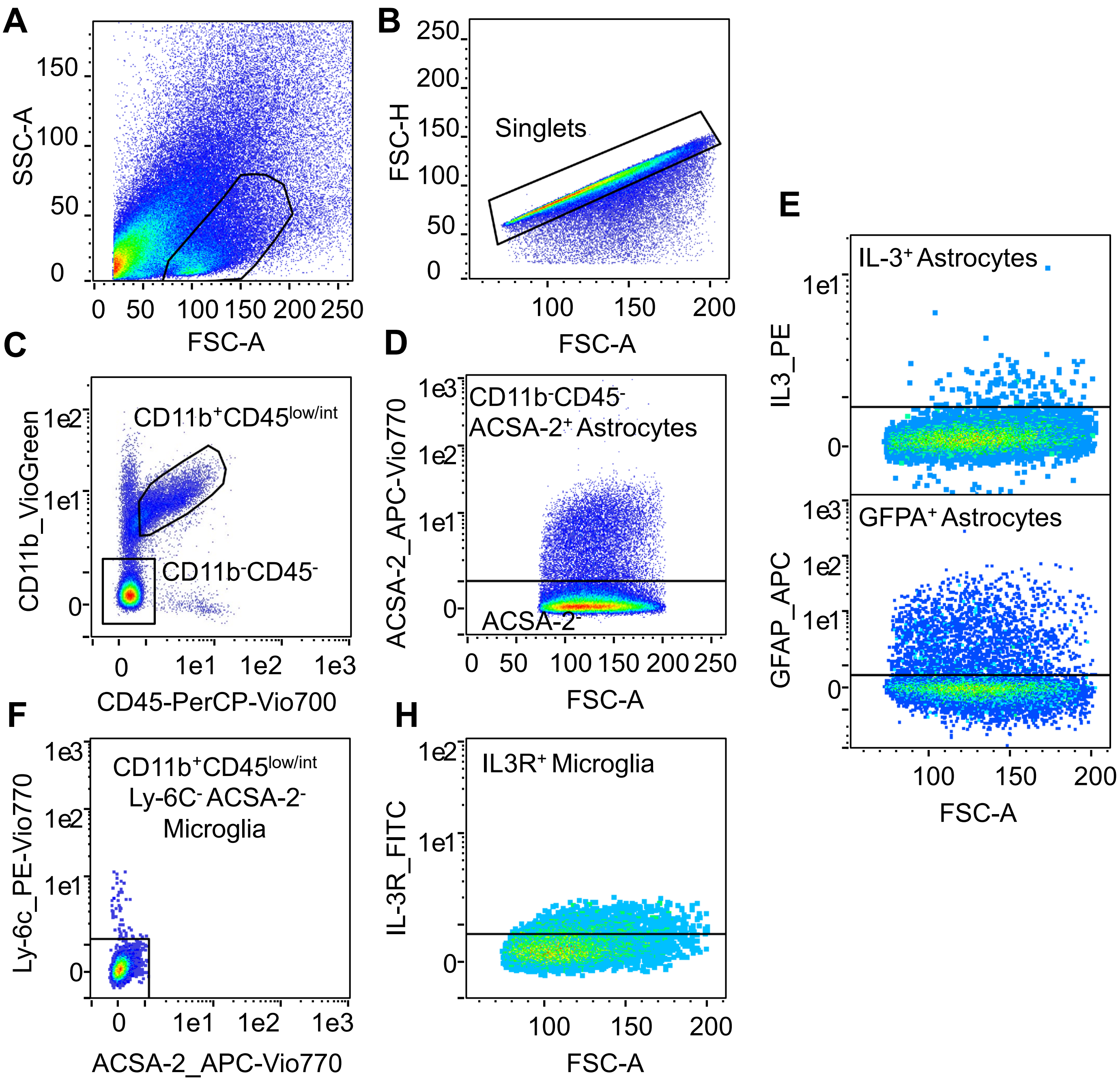


**Supplementary Figure 3. Gating strategy for IL-3/IL-3R analysis in astrocytes and microglia.** Single-cell suspensions from adult mouse brain were analyzed after fixation. Cells were first gated on (A) FSC-A versus SSC-A to exclude debris, (B) followed by FSC-A/FSC-H gating to select singlets. (C) From singlets, CD11b and CD45 expression were used to separate myeloid($CD11b^{+}CD45^{low/int}$) and non-myeloid populations ($CD11b^{-}CD45^{-}$). (D) The $CD11b^{-}$ $CD45^{-}$ population was used to identify astrocytes based on ACSA-2 expression, and (E) IL-3 and GFAP expression were assessed within the ACSA-2$^{+}$ astrocyte population (D). In parallel, the CD11b$^{+}$ CD45^low/int population (C) was further analyzed by (F) Ly-6C and ACSA-2 expression to define microglia as CD11b$^{+}$ CD45^low/int Ly-6C$^{-}$ ACSA-2$^{-}$ cells. (H) IL-3 receptor (IL-3R) expression was subsequently assessed within the gated microglia population.

#### Supplementary Figure 4

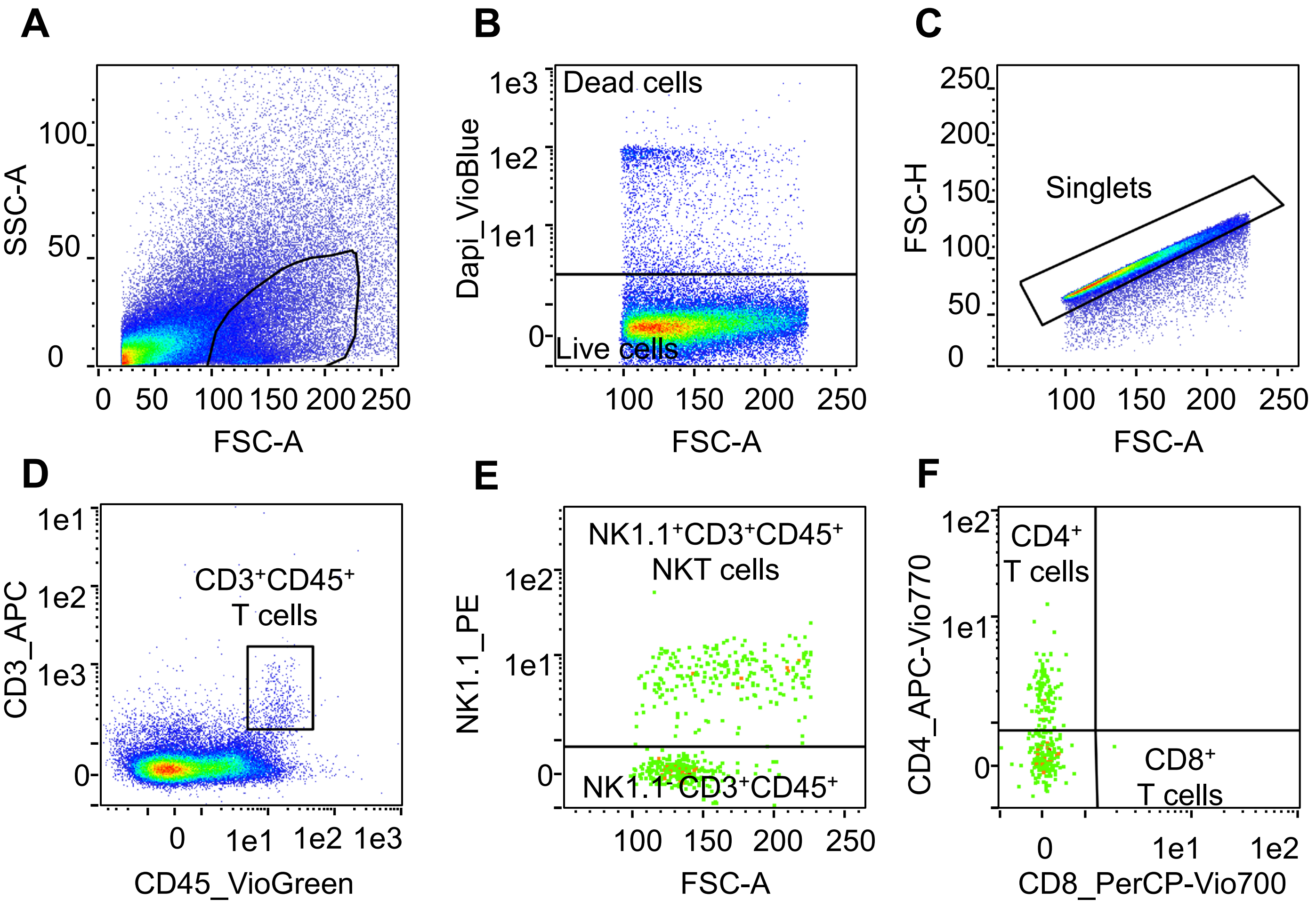


**Supplementary Figure 4. Flow cytometric gating strategy for identification of T cell and NKT cell populations in brain and meninges.** Single-cell suspensions from mouse brain and meninges were analyzed using the same gating strategy; representative plots shown are from brain. (A) Cells were first gated on FSC-A versus SSC-A to exclude debris. (B) Dead cells were excluded using a viability dye to identify live cells, (C) followed by FSC-A/FSC-H gating to select singlets. (D) From live singlets, CD45 and CD3 expression was used to define T cells ($CD3^+$ $CD45^+$). (E) Within the $CD3^+$ $CD45^+$ population, NKT cells were identified based on NK1.1 expression ($NK1.1^+$ $CD3^+$ $CD45^+$), while $NK1.1^-$ $CD3^+$ $CD45^+$ cells were classified as (F) $CD4^+$ and $CD8^+$ T cell subsets

### Supplementary Figure 5

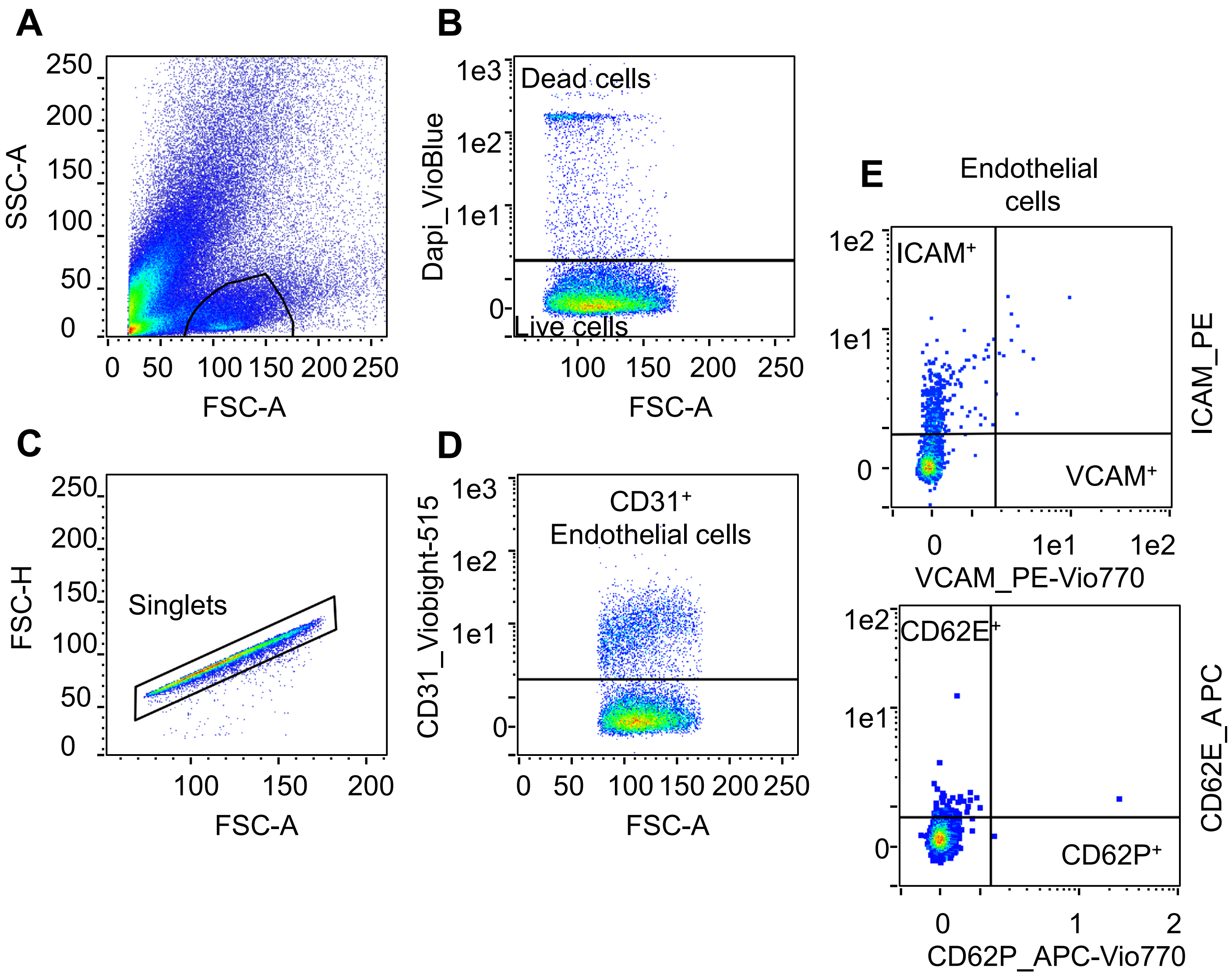


**Supplementary Figure 5. Flow cytometric gating strategy for identification of activated endothelial cells in the brain.** (A) Single-cell suspensions from mouse brain were first gated on FSC-A versus SSC-A to exclude debris. (B) Dead cells were excluded using a viability dye to identify live cells. (C) FSC-A versus FSC-H gating was used to select singlets. (D) From live singlets, endothelial cells were identified based on CD31 expression. (E) Within the $CD31^+$ population, activated endothelial cells were identified by expression of ICAM-1 ($ICAM\text{-}1^+$ $CD31^+$), VCAM-1 ($VCAM\text{-}1^+$ $CD31^+$), E-selectin ($CD62E^+$ $CD31^+$), and P-selectin ($CD62P^+$ $CD31^+$).
